# Evolution of histone mRNA 3’ end processing machinery in eukaryotes

**DOI:** 10.64898/2026.10.14.759114

**Authors:** Bin Zhao, Ursula Bond, Máire Ní Leathlobhair

## Abstract

Replication-dependent histone mRNAs in metazoans terminate in a conserved 3′ stem-loop (SL) recognized by stem-loop binding protein (SLBP) and processed by a specialized U7 snRNP-dependent cleavage machinery. Outside animals, the evolutionary history of this pathway remains poorly resolved, and fungi and most plants are widely thought to have lost SLBP-SL-dependent histone mRNA processing entirely. Here, we combine large-scale homology searches, structural and domain analyses, and genome-wide mapping of histone 3′-end features across eukaryotes to reconstruct the evolution of histone mRNA processing machinery.

We show that SLBP, stem-loop motifs and multiple U7-associated factors are far more broadly distributed across eukaryotes than previously appreciated, revising earlier models of widespread loss outside metazoans. Contrary to prevailing views, early-diverging fungal lineages retain SLBP orthologs with conserved RNA-binding domains, stem-loop motifs and additional components of the canonical pathway. Across fungi, we identify a clade-wise shift from SL-associated processing features toward canonical polyadenylation-associated features. Early-diverging fungi retain noncanonical poly(A) polymerases alongside SLBP and stem-loop motifs despite sparse U7-associated machinery, consistent with a previously unrecognized intermediate state in histone mRNA 3′-end processing. Our results support a stepwise evolutionary transition from ancestral SLBP/stem-loop-associated histone mRNA processing to canonical polyadenylation and provide a revised framework for the evolution of histone mRNA 3′-end formation in eukaryotes.

## Introduction

Replication-dependent (RD) histone messenger RNAs (mRNAs) terminate in a conserved 3’ stem-loop (SL) instead of the poly(A) tail found at the 3’ end of most eukaryotic mRNAs. An early step in histone mRNA 3’ end formation involves the binding of a stem-loop binding protein (SLBP) to the stem-loop structure. SLBP specifically recognizes the stem-loop to regulate histone mRNA metabolism, and the histone 3’-5’ exonuclease 3’hExo trims its 3’ end after processing (**Fig. 1A**)^1–4^. As well as recognizing the SL RNA, SLBP also has a role in the translation, export, and stability of mature histone mRNAs^5–8^.

**Figure 1.**
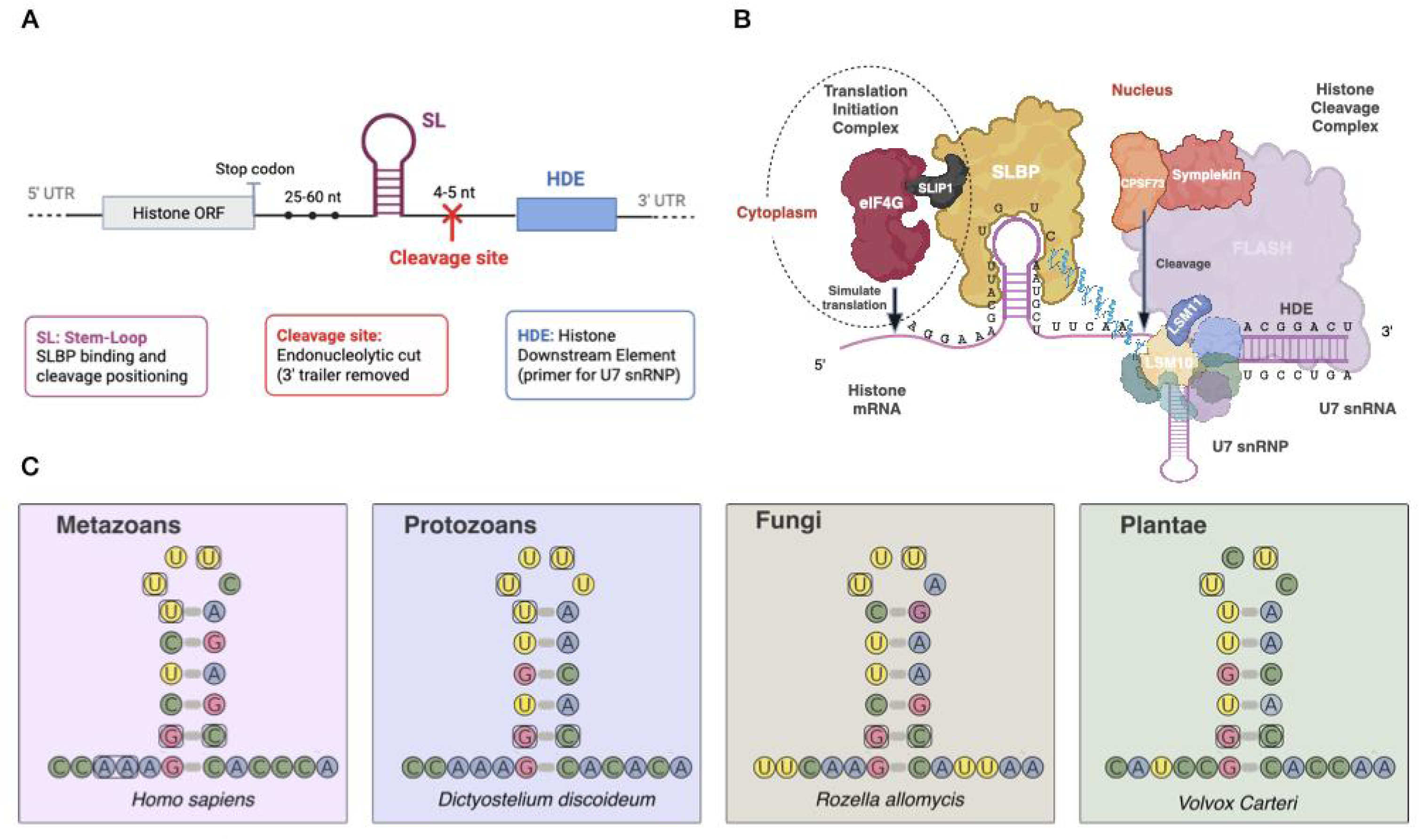
**(A) Structure.** The stem-loop of a canonical histone mRNA consists of a six–base pair stem and a four-base loop, as well as flanking sequences at both ends. The mRNAs have short 5′ and 3’ UTRs. The distance from the stop codon to the start of the conserved 26 nucleotide (nt) sequence at the 3’ end of the mRNA is 25–60 nucleotides and this distance is critical for regulation of histone mRNA stability. (**B) Processing**. These pre-mRNAs contain the conserved stem–loop sequence that binds stem–loop binding protein (SLBP) followed by the histone downstream element (HDE), which base-pairs with U7 small nuclear RNA (snRNA). A cleavage complex containing CPSF73 (cleavage and polyadenylation specificity factor subunit 73), symplekin, and other factors (not shown) is recruited to cleave the pre-mRNA. Within this complex, CPSF73 is the endonuclease that performs the cleavage, which occurs five nucleotides downstream of the stem–loop and upstream of the HDE. The U7 snRNA is a component of the U7 small nuclear ribonucleoprotein (snRNP), which contains a heptameric ring of five Sm proteins and two U7 snRNP-specific Sm-like proteins, LSM11 and LSM10. LSM11 contacts a 100 kDa zinc finger protein (ZFP100), which also interacts with the SLBP-stem-loop complex. **(C) Stem-loop conservation.** Representative histone mRNA stem-loop structures from a metazoan, a protozoan, a fungus, and a plant illustrate the conserved features of the canonical histone 3’ SL motif across major eukaryotic lineages. Despite substantial divergence, all four structures retain a conserved GC base pair at the second position of the stem and a conserved U at the third position of the loop.

SLBP functions together with the Histone Cleavage Complex (HCC), which includes the U7 small nuclear ribonucleoprotein (snRNP), FLICE-associated huge protein (FLASH), Symplekin, the endonuclease CPSF73, with additional cleavage/polyadenylation factors shared with canonical cleavage and polyadenylation (CPA) pathways^9–18^. The U7 snRNP binds directly to the histone downstream element (HDE) through base pairing between a purine-rich sequence at the 5′ end of the U7 snRNA and the histone pre-mRNA^19,20^. SLBP associates with the HCC through interaction with FLASH and a zinc finger protein ZFP100 (also known as ZNF473)^10,21,22^. This canonical SLBP-U7-dependent pathway (**Fig. 1B**) produces non-polyadenylated histone mRNAs that are rapidly degraded at the end of S phase ^6^.

Comparative analyses suggest that components of the SLBP-U7-CPSF73 pathway arose early in eukaryotic evolution. SLBP homologs have been identified in several unicellular eukaryotes, including *Dictyostelium* and *Chlamydomonas*^23^. These organisms have stem-loops proximal to histone mRNA 3’ ends, consistent with an early evolutionary origin of SL-mediated regulation. At the same time, López and Samuelsson (2008) reported that fungi, most land plants and several protozoa appeared to lack recognizable SLBP homologs and stem loop structures. In species lacking SLBP, histone mRNAs tend to be processed by canonical poly(A) signal (PAS)-mediated cleavage and polyadenylation (PAS/CPA)^24^. These data led to a widely cited model^25–28^ proposing that SLBP/U7-dependent, non-polyadenylated histone 3’ ends are largely restricted to animals (with scattered unicellular representatives), whereas fungi and most plants either never acquired this machinery or lost it and reverted to PAS/CPA processing.

Previous analyses suggested that SLBP and stem-loop structures are broadly absent from fungi and most land plants^23^, based on the comparatively limited number of eukaryotic genomes and proteomes available at the time. Since then, genome sequencing and protein annotation resources have expanded substantially. Moreover, the reported distribution implies repeated and complete loss of a highly conserved histone mRNA processing mechanism outside metazoans. Motivated by both the increased availability of genomic data and the apparent non-parsimony of this pattern, we revisited the evolutionary distribution of histone mRNA 3′-end processing machinery by examining the phylogenetic distribution of SLBP and associated pathway components.

Here we present genomic, transcriptomic, and structural analyses providing evidence that fungi retain relics of ancestral SLBP-related RNA-binding modules linked to polyadenylation-competent pathways lacking a recognizable U7 snRNP. This combination of features is consistent with an intermediate evolutionary state between canonical SLBP-SL processing and modern polyadenylation. We identify fungal SLBP-like proteins containing homologous SLBP RNA-binding domains (RBD) and show that early-diverging fungal lineages preserve key stem-loop features described in metazoans, including the conserved basal G-C pair at the second position of the stem base, and a uridine at the third position of the loop (**Fig. 1C**), for SLBP-binding^29,30^. We further identify fungal GLD-2–like nucleotidyltransferases containing catalytic motifs associated with adenosine preference^31,32^. The co-occurrence of SLBP, conserved histone stem-loops and polyadenylation-associated machinery, despite sparse U7-associated factors, is consistent with a previously unrecognized intermediate state between canonical SLBP-SL-U7 processing and predominantly polyadenylated histone mRNAs. This configuration parallels aspects of noncanonical histone mRNA processing described during *Drosophila* oogenesis^33^.

Our results refine the evolutionary history of histone mRNA 3’-end formation and identify early-diverging fungi as candidate intermediates linking SLBP–SL–U7-dependent processing with poly(A)-dependent cleavage and polyadenylation, supporting a model of progressive transition between these RNA-processing mechanisms.

## Results

### A Detailed Survey of SL Motifs and SLBPs in Eukaryotes

To start, we systematically assessed the distribution of SLBP across eukaryotes using homology and domain detection strategies (see **Methods**). Using sequence similarity searches (BLASTP), we identified 51,458 candidate SLBP homologs across 4,252 species of metazoans, plants, protists and fungi, after filtering (**Supplementary Figure S2**). To capture more divergent homologs that BLASTP might miss, we combined these hits with additional candidates from TBLASTN searches, InterPro, and OrthoDB curations, yielding the total candidate pool to 56,519 sequences (**Supplementary Tables S2-S3**). SLBP homologs are characterized by a centrally located RNA-binding domain (RBD) that interacts with the histone mRNA stem-loop structure. Within this combined candidate set, we detected the RBD in 20% of sequences, confirming 11,232 sequences from 2,968 species as high-confidence SLBP homologs. A targeted BLASTP search using the human SLBP RBD as a query recovered 11 additional homologs that were not captured in the initial screen. Across the combined results from all search approaches, SLBP-like proteins were identified in several protozoa, including the deeply branching lineages *Monocercomonoides* and *Euglenozoa* (**Supplementary File 4**). Although many of these SL sequences are highly diverged at the nucleotide level (**Fig. 1C**), they retain the core structural features required for SLBP recognition.

This updated dataset revises long-standing assumptions about the evolutionary history of replication-dependent histone mRNA processing. Earlier work proposed repeated, independent loss of SLBP in several eukaryotic lineages. Our expanded genome-wide survey instead recovered complete SLBP homologs in multiple clades previously reported to lack them entirely. While SLBP is consistently absent from *Dikarya* (modern multicellular fungi and yeast lineages), we confirmed its presence in early-diverging fungal lineages such as Chytridiomycota, supporting a model of secondary loss. Similarly, we recovered SLBP homologs in seed plants as well as in green and red algae and mosses, confirming and extending earlier findings reported by López and Samuelsson (2008).

Across all protists examined, ∼5% of genera (106 of 2,138) retained both SLBP homologs and detectable SL motifs downstream of canonical histone genes, indicating that SL-dependent processing is more broadly distributed across eukaryotes than previously recognized (**Fig. 2**). In contrast, and despite targeted searches of whole genome sequence data (**Methods**), no SLBP-like sequences were detected in *Oxytricha*, *Paramecium*, *Hemimastigophora* or *Provora*, consistent with absence of detectable SLBP in these clades.

**Figure 2.**
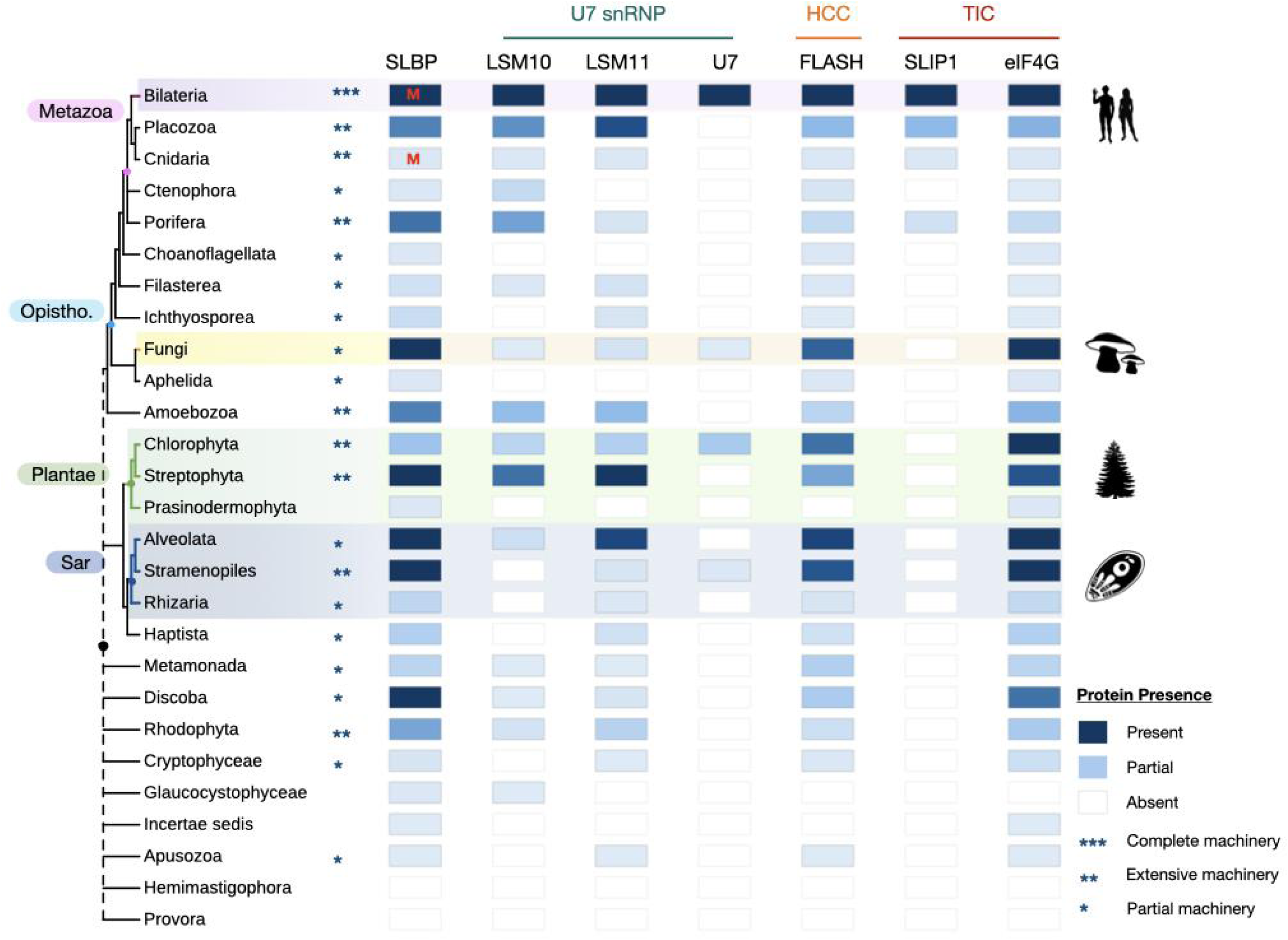
Distribution. Presence and relative abundance of canonical histone mRNA 3’-end processing factors (SLBP, SLIP1, U7 snRNP components and associated factors) across the eukaryotic taxa sampled in this study. Each square summarises the number of species within a clade in which a given factor was detected, with colour intensity proportional to the number of species encoding that factor (darkest shading, ≥100 species). The letter “M” within a square indicates that an SLBP N-terminal SLIP1-interacting motif was detected in that clade. Asterisks summarize the maximum complexity of the canonical processing machinery observed within individual species of each clade: *** complete machinery (all factors detected in one species); ** extensive machinery (≥ 5 factors detected in one species); and * partial machinery (≥ 3 factors detected in one species). Dashed branches indicate unresolved phylogenetic relationships. Silhouettes are adapted from www.phylopic.org. The eukaryotic tree is based on the NCBI Taxonomy Database and annotated using ITOL ^93^. HCC, histone cleavage complex.

The N-terminus of metazoan SLBP contains a short SLIP1-binding motif required for recruitment of SLIP1 and activation of replication-dependent histone mRNA translation. We therefore systematically searched for this motif in all SLBP orthologs (see **Methods**). This motif was readily detected in metazoan SLBPs (**Fig. 2**) and showed strong conservation of key residues (**Supplementary Figure S3**). Outside Metazoa, however, we did not detect convincing SLIP1-interacting motifs in any surveyed SLBP, including those from fungi, plants and unicellular eukaryotes. These results suggest that while SLBP and SL-mediated processing are retained in multiple non-metazoan lineages, the specific SLIP1-dependent translation activation mechanism is likely restricted to metazoans.

### A Survey of SLBPs outside Eukaryotes

Having identified widespread retention of SLBP-associated histone processing features across eukaryotes, we asked whether more distant homologs of the SLBP RNA-binding domain could be detected outside the eukaryotic tree. Sequence-based searches identified only two SLBP-like candidates in Archaea: one from an unclassified Methanobacteriota species (NCBI accession: RYG41222.1) and another from an undefined archaeal genome (UniParc: UPI000C600A47).

Similarly, three bacterial candidates were found to contain a detectable RBD. These included a *Legionellales bacterium* (NCBI accession: OGV26409.1), a *Crocinitomicaceae bacterium* (UniParc accession: UPI000C6245EF), and an *Optitutia bacterium* (UniParc accession: UPI000C5FD937). These bacteria belong to the phylum Pseudomonadota (formerly Proteobacteria), the Fibrobacteres-Chlorobi-Bacteriodetes (FCB) group, and the Planctomycetes-Cerrucomicrobia-Chlamydiae (PVC) superphylum, respectively. Despite structural similarity to the eukaryotic SLBP RNA-binding domain, these bacterial proteins generally lacked the conserved residues associated with recognition of the histone mRNA 3’ stem-loop, suggesting that they likely perform distinct RNA-binding functions. However, notably, in all bacteria where SLBP-like domains were predicted, the corresponding species, or closely related taxa, had previously been reported to encode histone-fold domain-containing proteins ^89,90^.

We also examined large double-stranded DNA viruses, including giant viruses reported to encode histone-like proteins and other chromatin-associated factors ^91,92^. However, neither sequence-based nor structure-based searches detected SLBP homologues or other canonical components of the histone 3’-end processing machinery in viral genomes.

### Eukaryotic Distribution of Histone 3’-End Processing Factors Beyond SLBP

To determine whether the broad distribution of SLBP extends to its processing and translation partners, we surveyed the U7 snRNP-specific Sm-like proteins LSM10 and LSM11, the scaffolding factor FLASH, and the translation factors SLIP1 and eIF4G (**Fig. 2**).

Homologs of both LSM10 and LSM11 were detected widely outside animals, including in green algae, land plants, red algae, stramenopiles, unicellular holozoans and amoebozoans. Examples included the brown alga *Ectocarpus* and the slime molds *Acytostelium subglobosum*, *Synstelium polycarpum*, and *Dictyostelium garhantuum*.

In fungi, however, we did not observe LSM10 and LSM11 together within the same species genome. LSM10 was only detected in the entomopathogenic fungus *Massospora platypediae*, whereas LSM11 was found in early-diverging fungal lineages including *Rozella allomycis* and the microsporidian *Nematocida displodere*. Thus, fungal retention of individual U7-associated proteins contrasts with the broader co-occurrence of LSM10 and LSM11 outside fungi.

FLASH (CASP8AP2), which links the SLBP-SL complex to the histone cleavage complex, (HCC), was also detected widely outside metazoans. Although full-length FLASH proteins were not detected in most fungi, proteins containing conserved FLASH-associated domains were identified in early-diverging fungi, including *R. allomycis* (**Supplementary Figure S5**). In *R. allomycis*, these candidates co-occurred with LSM11 and a predicted U7-like snRNA (**Fig. 2; Supplementary Figure S6**). These data suggest that functional components of the histone 3’-end cleavage machinery, including the FLASH-LSM11 interaction that stabilises U7 recruitment to histone pre-mRNAs, were likely present in early-branching opisthokonts, although it does not establish a functional FLASH–LSM11 interaction or a complete cleavage complex.

Following cleavage at the histone 3’-end, SLBP remains bound to the mature histone mRNA and promotes its translation. In metazoans, SLBP recruits SLIP1, which in turn cooperates with eIF4G to facilitate translation initiation. We identified eIF4G in several non-metazoan species encoding SLBP, including the green alga *Micromonas commoda*, the moss *Sphagnum magellanicum*, and in *R. allomycis*.

Canonical SLIP1, however, was confidently detected only in metazoans. A fragmented SLIP1-like sequence was identified in the red alga *Rhodymenia pacifica*, but no intact functional protein could be inferred. We recovered several SLIP1-like proteins with SLIP1-type HEAT-like repeat architectures but lacking one of the short SLBP-interacting motifs (**Supplementary Figure S3**). Together with the absence of detectable canonical SLIP1-binding motifs in non-metazoan SLBPs (**Fig. 2**), these findings indicate that the characterised SLBP-SLIP1 translation module has a narrower distribution than SLBP itself. The data therefore distinguish broad conservation of the RNA-binding component from conservation of its metazoan translation partners, though without resolving whether the specialised interaction arose in animals or was lost elsewhere.

### Conserved Functional Features of Eukaryotic stem-loop Histone mRNAs

To examine conservation and loss of stem-loop-mediated histone mRNA 3’-end regulation across eukaryotes, we analysed mRNAs encoding the four core histones (H2A, H2B, H3 and H4) across metazoans, protists, fungi and plants. For each coding sequence, we extracted downstream genomic flanking regions, identified candidate SL structures using a combined sequence-structure pipeline (**Methods**), and quantified SL sequence conservation and divergence, SL distance from the stop codon, and the presence of a downstream purine-rich HDE (**Fig. 3**).

**Figure 3.**
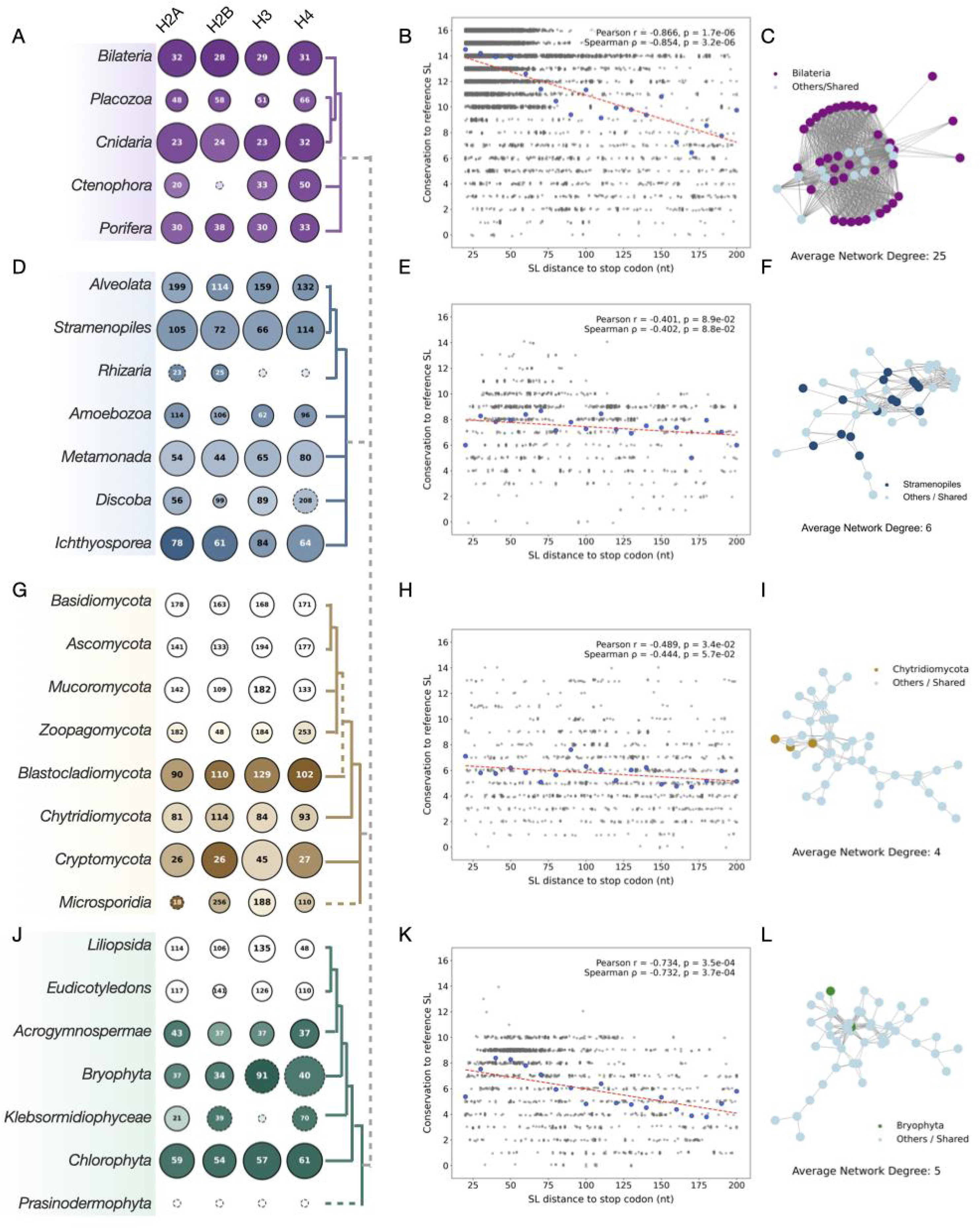
Phylogenetic distribution, conservation, and diversity of histone mRNA SL motifs across eukaryotes. **(A, D, G, J) Distribution**. Phylogenetic trees of metazoan, protozoan, fungal and plant lineages showing the distribution of SL motifs across core histones (H2A, H2B, H3 and H4). Each circle represents a clade-histone combination, with circle size indicating the relative abundance of SL-containing sequences normalized by the number of species in each clade (number of SLs divided by number of species). White circles indicate the absence of SLBP. Solid and dashed edges show the presence or absence of HDE in this lineage, respectively. Darker colours indicate greater SL sequence conservation within the group. Numbers within circles denote the median SL to stop codon distance. Phylogenetic relationships are based on the NCBI Taxonomy database. **(B,E,H,K) Correlation**. The relationship of SL conservation to stop codon. Grey points represent individual SLs, blue points indicate binned means (10-nt intervals), and dashed red lines show linear regression fits with Pearson’s r and Spearman’s rho values. **(C, F, I, L) Similarity**. In the SL similarity network of the 50 most frequent SL core variants within each group, nodes represent distinct SL 16-bp cores (sharing the same MFE, delta kcal/mol) connected by edges reflecting sequence similarity. Node colours show representative clades distribution.

### Strong conservation of histone stem-loop sequence and positioning in metazoans

Across 2,281 metazoan species, SL motifs were consistently positioned close to the coding region, with a median distance of 30 nt downstream of the stop codon (**Fig. 3A**). In representative model organisms (*Homo sapiens*, *Mus musculus*, *Drosophila melanogaster* and *Danio rerio*), the diagnostic G-C base pair at stem position 2 and the conserved uridine at loop position 3 were present in ∼95–100% of histone mRNAs, and SLs were typically located 23-31 nt downstream. SLs were generally followed by purine-rich HDE-like regions.

SL conservation extended across non-bilaterian metazoans, including cnidarians, placozoans, ctenophores and sponges, in which stem loops were detected downstream of core histone genes with complete conservation of the key G-C stem pair and stem loop U and relatively constrained distances from stop codons (**Fig. 3A; Supplementary Table S10**). All identified SLs were followed by strong purine-rich HDEs, except for H4 in the cnidarian *Clytia hemisphaerica* due to insufficient length of flanking sequence.

We next examined the diversity of SL sequence variants and their relationships using a similarity-network representation of the 16-nt SL core (**Methods**). Across all high-confidence metazoan SLs (n = 37,587), SL hairpins showed high sequence conservation, with an average of 14 of 16 positions identical to the reference stem loop (**Methods**). SL conservation dropped sharply as SL distance from the stop codon increased (Pearson r = -0.87; Spearman rho = -0.85; both p < 1e-5; **Fig. 3B**). Sequence similarity networks across all unique metazoan SL sequences revealed a densely connected topology (**Fig. 3C**), with predicted folding stabilities distributed between -7 and -10 kcal/mol minimum free energy (MFE). The top 50 most frequent SL variants formed a robust core network with an average network degree of 25. Together, these results indicate strong evolutionary constraint on SL sequence, structure and placement across metazoans.

### Partial conservation of the SL-SLBP system in protists

In protists, SL conservation was more variable than in metazoans, but remained widespread (**Fig. 3D**). Of 310 surveyed protozoan species, 214 encoded detectable SL structures downstream of histone genes, frequently in lineages that also encoded SLBP (**Fig. 3D**). In several groups, including Ichthyosporea and Rhizaria, stem loops retained diagnostic positions in all core histones, were located near stop codons, and SLBP was present, whereas in Amoebozoa, Stramenopiles and other protist lineages SLs were typically positioned further downstream and showed reduced sequence conservation (**Supplementary Table S10**).

Across all protist SLs (n = 1,109 after filtering), the association between SL distance and sequence conservation was weaker and not statistically significant (Pearson r = -0.4; Spearman rho = - 0.4; **Fig. 3E**). SL similarity sequence networks were fragmented, and the top 50 most frequent SLs formed a weakly connected network (**Fig. 3F**), with an average degree of 6. Predicted folding stabilities ranged broadly from unstable (∼0 kcal/mol) to moderately stable (-8 kcal/mol). These results indicate that while many protists retain SLBP-associated stem-loop features, their structure and positioning have diversified substantially relative to metazoans.

### Divergence and partial decay of the SLBP-SL processing system across fungal phyla

In fungi, SLs were present but often restricted to subsets of histone genes and showed substantial variability in both sequence conservation and position relative to the stop codon. Cryptomycota retained comparatively conserved SL features. In *R. allomycis*, SLs were detected downstream of all core histones and were typically positioned ∼26 nt from the stop codon, whereas *Paramicrosporidium saccamoebae* retained SLs downstream of most histones but lacked a detectable SL for H3. Chytridiomycota showed much greater positional variability and partial retention of SL motifs across histone classes. In some Chytridiomycetes, SLs were retained only downstream of individual histone genes, whereas other loci lacked detectable motifs entirely. Blastocladiomycota exhibited narrower SL distance distributions and moderately conserved motifs (**Fig. 3G**). Across these three phyla, no consistent restriction of SL retention to a certain histone type was observed.

An SLBP ortholog was detected in Zoopagomycota (*Coemansia spiralis*), but no SL motifs could be identified downstream of histone genes. Conversely, in Mucoromycota, Ascomycota and Basidiomycota SL-like hairpins were detectable, despite the apparent absence of SLBP orthologs.

Microsporidia represented the most extreme case. SL-like structures were located far downstream (∼190 nt), conservation of diagnostic SL positions was low (20% G-C at stem position 2; 36% loop U; **Fig. 3G**), and HDE-like purine-rich regions were rarely detected. In most species, SL motifs were only detectable primarily downstream of H3 loci.

Fungal SL candidates showed low sequence identity to the metazoan reference overall (average 5.8/16 positions; n = 1,177 after filtering). The association between SL distance and sequence conservation was moderate but not statistically significant (Pearson r = -0.49; Spearman rho = - 0.44; **Fig. 3H**). Stem-loop similarity networks were fragmented and weakly connected, and predicted folding energies ranged from −10 to 0 kcal/mol (**Fig. 3I**). SL retention and conservation varied both between and within fungal lineages, ranging from comparatively conserved motifs in early-diverging lineages to minimal detectable conservation in Microsporidia, with SL distance and sequence conservation only weakly associated across this lineage.

### Retention, remodeling and decoupling of SLBP and SL in plants

In plants, SL-associated histone processing features were retained but showed increasing divergence across the lineage (**Fig. 3J**). Green algae and early-diverging land plants retained relatively canonical SL features, including SLs positioned close to stop codons and strong conservation of diagnostic positions associated with SLBP recognition. In chlorophytes such as *Chlamydomonas reinhardtii* and *Volvox carteri*, SLs were detected downstream of all four core histone genes together with purine-rich HDE elements (**Fig. 3J**), whereas bryophytes and gymnosperms also retained strong conservation of the characteristic stem G-C pair and loop uridine.

By contrast, later-diverging streptophyte algae and monocot flowering plants (Liliopsida) frequently contained more divergent SL-like hairpins, with weaker conservation of diagnostic positions and greater positional variability relative to the stop codon. In Klebsormidium nitens, for example, SLs were detected downstream of H2A, H2B and H4 but not H3, and the canonical loop uridine was absent (**Fig. 3J**). Across plant SLs (n = 1,570), sequence conservation was intermediate overall (mean 6.7/16 identical with reference SL), and conservation decreased with increasing distance from the stop codon (Pearson r = -0.43 and Spearman rho = -0.43; both p < 0.005; **Fig. 3K**). SL similarity networks showed moderate connectivity (average degree, 5; **Fig. 3L**), and varying predicted folding energies (-1 to -8 kcal/mol).

### Stem-loop and polyadenylation-associated features coexist in early-diverging fungi

To assess how histone mRNA 3’-end processing varies across early-diverging fungi, we classified histone loci based on regulatory elements detected in their downstream regions. Specifically, we tested whether histone transcripts retained stem-loop features associated with SLBP-dependent processing and canonical cleavage/polyadenylation. Loci were assigned to one of four states: SLBP-SL, SLBP-SL-PAS, SLBP-PAS, or PAS-only (see **Methods**). We then quantified the relative prevalence of these states within species and across fungal clades.

Among loci that could be assigned to one of the four defined processing types, the distributions support a progressive shift from SL-associated histone 3′-end processing towards canonical polyadenylation (**Fig. 4A-C**). Cryptomycota and Blastocladiomycota were dominated by loci containing both SL-associated features and PAS elements (SLBP-SL-PAS), and the clade-level contrast between SL-associated and PAS-only states was strongly positive (**Fig. 4C**), indicating overall enrichment of SL-associated processing features in these groups.

**Figure 4.**
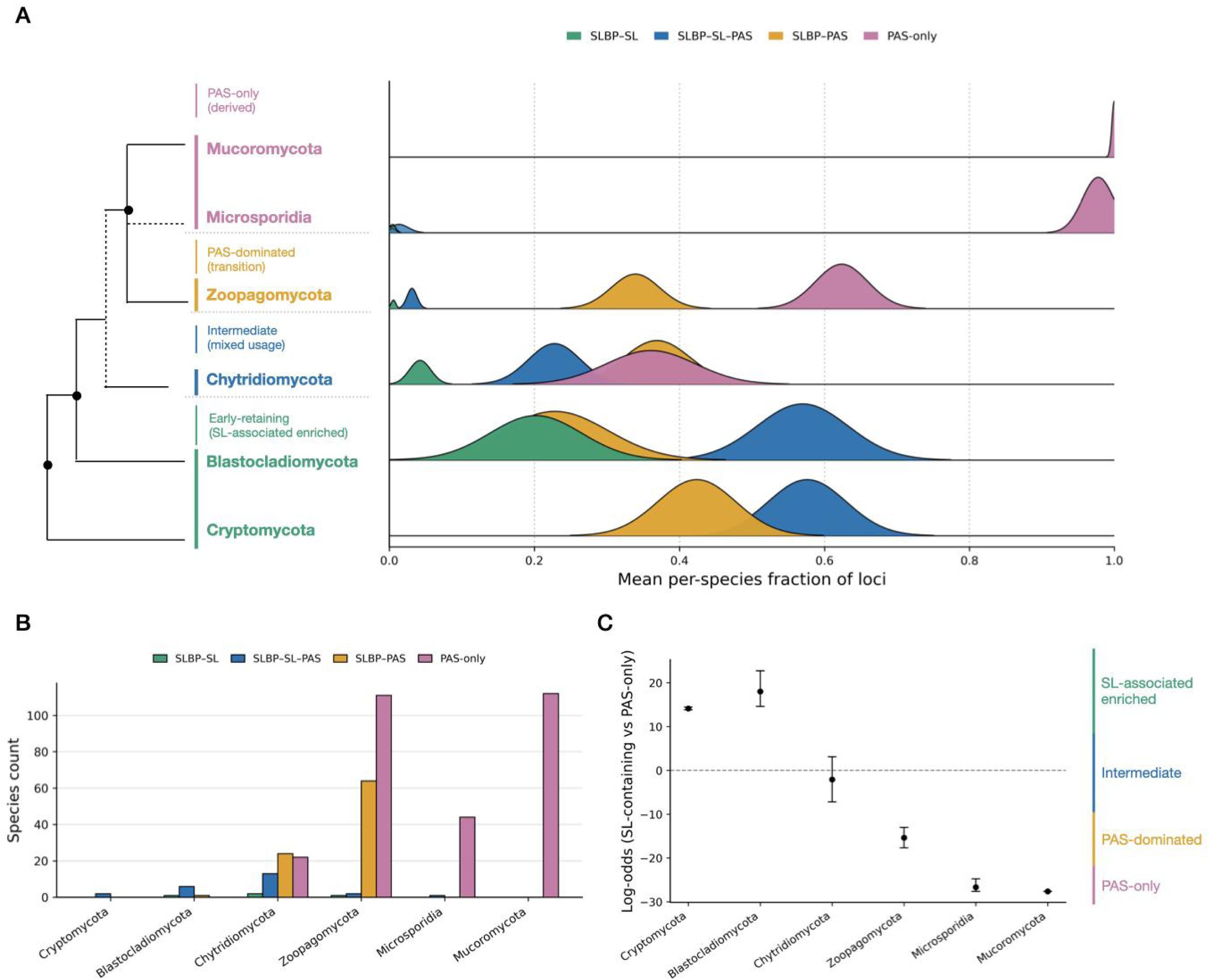
Histone 3’-end processing modes in early-diverging fungi. **(A) Species-level state distributions.** Ridgeline profiles summarize the four mutually exclusive processing states across species within each clade. For each state, horizontal position indicates the mean per-species fraction of evaluable histone loci assigned to that state (0-1), and profile width represents the standard error of this mean. Profile height encodes the fraction of species in the clade in which the state is present, square-root transformed for visualization. (**B) Dominant state per species.** Dominant state is defined as the state with the highest species-level fraction, with ties assigned to the coexistence class (SLBP-SL-PAS). Counts show the distribution of dominant states across clades. (**C) Clade-wise contrast of SL-containing vs PAS-only states.** Mean log-odds across species are shown with 95% bootstrap CIs (B = 3,000). Values >0 indicate enrichment of SL-containing states; values near 0 indicate similar representation; values <0 indicate enrichment of the PAS-only state

Chytridiomycota showed a heterogeneous distribution of histone 3′-end features. Clade means were partitioned across SLBP-SL-PAS, SLBP-PAS, and PAS-only classes; species-level fractions are also multi-modal (**Fig. 4A**), the dominant processing state is relatively equal/mixed (**Fig. 4B**), and the clade-level contrast is near zero (**Fig. 4C**). These data are consistent with erosion or replacement of SL-associated processing across different histone loci within the same genomes.

PAS-associated configurations predominated in Zoopagomycota, Mucoromycota and Microsporidia. Zoopagomycota contained primarily PAS-only loci alongside a smaller proportion of SLBP-PAS configurations, whereas classifiable loci in Mucoromycota and Microsporidia were almost exclusively PAS-only. Clade-level contrasts were strongly negative in these groups (**Fig. 4A-C**). These results are consistent with a progressive shift from coexistence in Cryptomycota and Blastocladiomycota, to heterogeneous intermediate states in Chytridiomycota, to PAS-dominated processing in Zoopagomycota, and to PAS-only processing in Mucoromycota and Microsporidia.

To evaluate whether these inferred processing states were reflected at the transcript level, we surveyed publicly available RNA-seq datasets from early-diverging fungal lineages and examined histone transcript termini where coverage permitted (**Supplementary Figure S8**). In Blastocladiomycota, *Allomyces macrogynus* H2A contains a PAS variant (AATATA) immediately upstream of a clear poly(A) tail and downstream of an SL structure, consistent with an SLBP-SL-PAS configuration. *Catenaria anguilulae* H4 likewise carries an SL followed by a poly(A) tail with no detectable PAS, consistent with coexistence of SL-associated and polyadenylated features at the locus level.

Chytridiomycota again showed marked within-genome heterogeneity, with different histone transcripts terminating through distinct combinations of SLs, canonical or variant PAS motifs and poly(A) tails (**Fig. 4B-C**). By contrast, zoopagomycete and mucoromycete histone transcripts predominantly terminate in PAS-associated poly(A) tails without detectable SLs, concordant with the PAS-dominated states inferred from genomic data (**Fig. 4C)**.

The locus-level patterns raised an important mechanistic question. Several early-diverging fungi retain SLBP and stem-loop motifs despite lacking most recognizable U7-associated factors, whereas multiple histone transcripts terminate in poly(A) tails despite lacking detectable canonical polyadenylation signals (**Supplementary Figure S8**). These observations were not readily explained by either the canonical metazoan SLBP-SL-U7 pathway or conventional PAS-directed cleavage and polyadenylation. One possibility is that histone transcript maturation in these lineages involves noncanonical poly(A) polymerases.

### Candidate alternative poly(A) polymerases are retained in SLBP-positive fungi

In *Drosophila* oocytes, histone transcripts can undergo SLBP-associated cytoplasmic polyadenylation mediated by the noncanonical poly(A) polymerase GLD2. Furthermore, in *Saccharomyces cerevisiae*, factors associated with transcription termination and 3’ end processing of small, non-coding RNAs, such as the Nab/Nrd/Sen^94^ and the nuclear TRAMP/exosome^95,96^ complexes, have been shown to play a role in regulating replication dependent histone mRNAs. *S. cerevisiae* histone mRNAs have shorter than average poly(A) tails and the tail length varies during the cell cycle: G1 cells have poly(A) tails of >50 nt while newly synthesised histone RNAs in S-phase have tails <25 nt^94^. These short poly(A) tails on newly replicated histone mRNAs are reminiscent of the very short poly(A) tails added by the TRAMP-associated non-canonical Trf4 poly(A) Polymerase during the 3’ end processing of small non-coding RNAs^97^. The coexistence of histone stem-loops and polyadenylated transcripts in lineages lacking recognizable U7 prompted us to ask which alternative poly(A) polymerases are retained in these organisms. We therefore surveyed fungal genomes for noncanonical poly(A) polymerases and compared their distributions with SLBP and U7-associated factors.

Our survey of histone mRNA processing proteins across fungal species (**Fig. 5A**) revealed that canonical metazoan histone 3’-end processing components are largely absent from fungi, whereas SLBP and noncanonical poly(A) polymerases (ncPAPs; classified here as GLD2-like and Trf4-like proteins) are retained together in several early-diverging lineages. GLD2-like candidates were comparatively restricted, being detected only in Chytridiomycota, Zoopagomycota, Mucoromycota and Dikarya. In contrast, Trf4-like ncPAP candidates were detected across all surveyed fungal lineages (**Fig. 5A**). Their broad distribution showed little correspondence with the retention of SLBP or histone SL features, arguing against Trf4-like proteins alone accounting for the inferred transition in histone mRNA processing.

**Figure 5.**
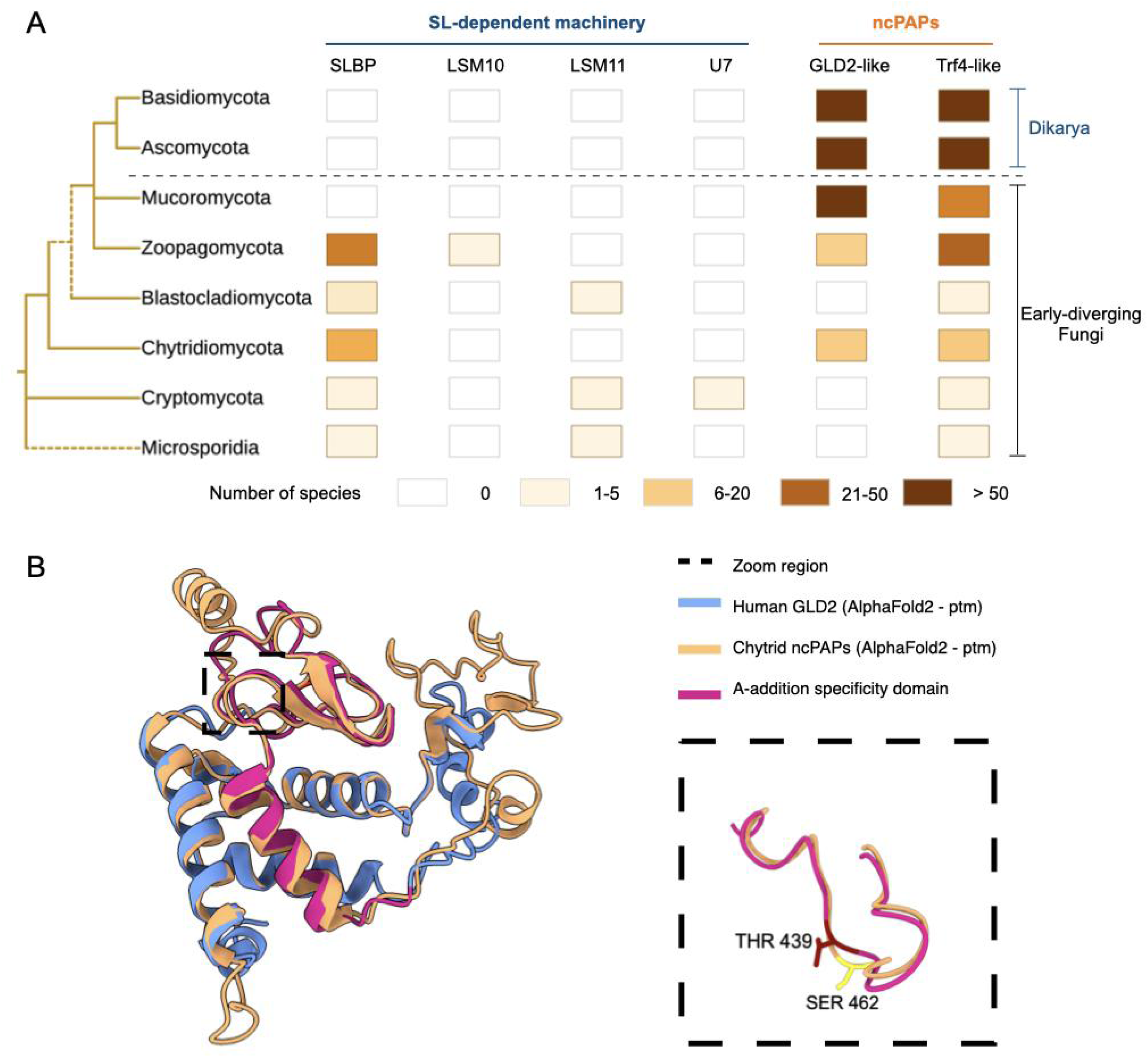
Fungal noncanonical poly(A) polymerases and their relationship to histone mRNA processing. **(A) Distribution.** Factors involved in fungal histone mRNA processing machinery. Each square represents the detection status of a given protein in a given clade. The first four columns correspond to the canonical SL-dependent histone mRNA 3’end processing complex. The last two groups are GLD2-like and Trf4-like proteins in non-canonical poly(A) polymerases (ncPAPs). Phylogenetic relationship is adopted from the MycoCosm database ^98^ (**B) Structural comparison.** Alphafold models of human GLD2 (Uniprot: Q6PIY7; blue) and a representative chytrid ncPAP from Cladochytrum replicatum (NCBI: KAI8803508.1; brown) aligned in UCSF ChimeraX. Only the region shared between both proteins is shown. The A-addition specificity domain of GLD2 (human residues 384-442) is highlighted in red. The inset shows the region surrounding human residue THR 439 and the corresponding residue in the chytrid protein, implicated in nucleotide preference in GLD2-family polymerases.

To assess whether candidate factors are expressed in early-diverging fungi, we quantified transcript abundance in publicly available RNA-seq datasets from zoospore and multinucleate stages of *Spizellomyces punctatus* (**Supplementary Table S9; Supplementary Figures S9**). Both SLBP and the candidate ncPAP were substantially enriched in multinucleate growth stages relative to zoospores, paralleling the expression of a PAS-lacking histone H2A locus and core histone genes. Thus, SLBP and a candidate alternative poly(A) polymerase are co-expressed in a developmental stage associated with increased histone expression, although these data do not establish a functional relationship between them.

Structural analysis further supported the annotation of these proteins as candidate ncPAPs. AlphaFold2 models of representative fungal ncPAPs showed strong structural conservation of the nucleotidyltransferase catalytic core (RMSD ∼1 Å) relative to human GLD2, including conservation of the PAP_assoc domain and the surrounding catalytic region (RMSD ∼1 Å) (**Fig. 5B**). The PAP_assoc loop aligns with the region surrounding the T439-N440 segment of human GLD2, and the corresponding site in fungal ncPAPs adopts the non-histidine residue that avoids switching ATP specificity to UTP. These structural features are compatible with fungal ncPAPs that preferentially incorporate adenosine and identify these proteins as candidates that could contribute to polyadenylation of fungal histone transcripts.

## Discussion

Our results revise the view that replication-dependent histone mRNA 3’-end processing is specific to metazoans and instead suggest that the underlying components of this pathway originated early in eukaryotic evolution and have subsequently been modified in different lineages (**Fig. 6A)**. By combining dense SLBP homology searches, systematic mapping of SL motifs, and comparative analysis of early-diverging fungi, we identify a continuum of genome configurations defined by SLBP binding sites, SL motifs, polyadenylation signals, and poly(A) polymerase. These configurations range from the canonical SLBP-SL-U7-dependent cleavage pathway to PAS-dependent cleavage and polyadenylation. Rather than a single loss of the SLBP-SL pathway in fungi, the data suggest instead a progressive decoupling among core components: the SLBP protein, SL hairpin motifs in histone transcripts, and PAS-mediated cleavage signals (**Fig. 6B)**.

**Figure 6.**
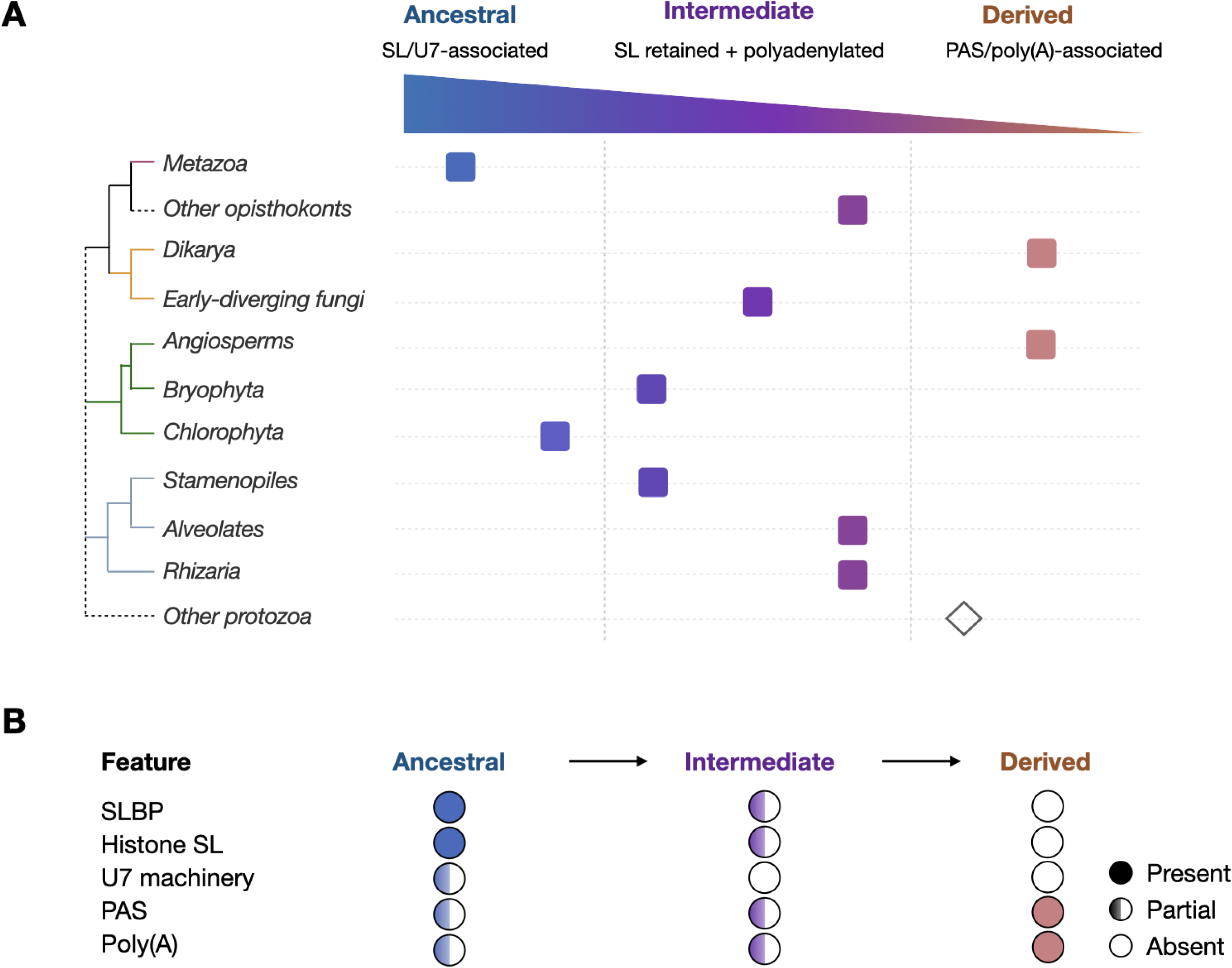
Evolutionary trend in histone mRNA 3’ end processing capacity across eukaryotes. **(A)** Phylogenetic summary. Coloured squares indicate clade level inferred processing capacity, and the open diamond denotes uncertain cases in other protozoan species. **(B)** Proposed Fungal transition model. An ancestral state retains SLBP, SL and U7 machinery, whereas the derived state is dominated by PAS/Poly(A)-associated processing. The intermediate state contains coexistence of both machinery factors but the U7 related factors are mostly lost. The filled, half-filled and open circles indicate present, partial and absent features, respectively.

Across fungi, the differing distributions of SLBP, histone stem-loop motifs and PAS elements suggest that these components have not been retained or lost as a single unit. PAS-associated configurations are more prevalent in some clades, whereas others retain diverse combinations of SL-associated and polyadenylation-associated features. Some loci retain SL-like hairpins alongside PAS elements or in species lacking detectable SLBP, while other species retain SLBP without a detectable canonical SL at particular histone loci. This variation supports a model of remodelling through decoupling of RNA recognition and downstream processing machinery. The greater distances between stop codons and candidate SLs observed in Zoopagomycota and Microsporidia are consistent with relaxed selection on these elements. In Microsporidia and some Mucoromycota, distant or weak SL-like structures, detectable PAS elements and the absence of detectable SLBP together suggest predominantly PAS-associated histone mRNA processing

Protein distributions in fungi support an intermediate state of histone mRNA processing characterized by SLBP retention and polyadenylation in the absence of recognizable U7. In several early-diverging lineages, SLBP orthologs and SL motifs co-occur with candidate noncanonical poly(A) polymerases, whereas no broadly conserved recognizable U7 system is detected. This distribution argues against conservation of a canonical metazoan-like U7-dependent cleavage complex^99^. The ncPAP families, however, show different evolutionary distributions. Trf4-like proteins are broadly distributed across fungi and show little correspondence with retention of SLBP or histone SL features. Because Trf4-family polymerases are principally associated with nuclear RNA surveillance, their presence alone does not identify them as histone mRNA-processing enzymes. By contrast, GLD2-like ncPAPs have a more restricted distribution that overlaps several SLBP-positive early-diverging lineages. The retention of GLD2-like ncPAPs identifies one candidate mechanism that can now be tested experimentally. A precedent for the coexistence of SLBP recognition and polyadenylation occurs during animal oogenesis, where SLBP participates in cytoplasmic polyadenylation of maternal histone mRNAs^33^. Fungal lineages retaining this combination of features may therefore represent evolutionary intermediates between canonical SLBP-SL-U7-dependent processing and PAS-directed cleavage and polyadenylation.

Genetic studies in *Drosophila* suggest why such intermediates may arise from an ancestral non-polyadenylated system. SLBP and U7 snRNP cooperate to generate the canonical histone 3’ end, and mutations in either SLBP, U7 snRNA, or in the U7-specific LSM10 and LSM11 disrupt histone pre-mRNA processing, yielding polyadenylated histone mRNAs and severe developmental defects ^55,100^. These data indicate that the fully non-polyadenylated histone mRNA pathway requires both SLBP and U7 snRNP. The emergence of U7 and its dedicated LSM10/11 proteins likely represents a critical step toward the specialized metazoan pathway, with the broader phylogenetic distribution of SLBP being consistent with an earlier origin of the RNA-recognition component^23,28^.

This proposed remodelling of SLBP is further supported by its sequence properties in fungi. Comparison of fungal SLBP orthologs with their bilaterian counterparts shows that fungal SLBPs lack the SLIP1-interacting motif required for SLIP1-dependent translational activation in metazoans, while the RBD remains well conserved (**Fig. 2**). This pattern is consistent with retention of the RNA-recognition function of SLBP in fungi alongside divergence of its associated translation interactions.

Plants show a related pattern. Histone SL sequences, predicted structural stability, and spacing relative to the stop codon have diversified across the lineage. This is consistent with partial retention of SLBP-SL processing. At the same time, the sequence and positional constraints normally required for canonical SLBP recognition have progressively relaxed. As in fungi, this points to remodelling of individual pathway components, not loss of the pathway as a whole. At the scale of the eukaryotic tree, our expanded SLBP survey and SL mapping refine earlier conclusions about the distribution of histone 3’-end processing^23^. We identify high-confidence SLBP orthologs with recognizable RBD in plants, algae, protists and multiple early-diverging opisthokont lineages, together with stem loops downstream of histone genes in many of the same taxa. Outside metazoans, these SLs are more diverse in sequence, position relative to the stop codon, and predicted stability, but often retain features compatible with SLBP recognition. The most parsimonious explanation of the data is that the SLBP protein and the stem-loop it recognizes in histone mRNAs originated early in eukaryotic evolution and were present in the last common eukaryotic ancestor. Subsequent lineages appear to have retained, modified or lost individual components of this module to different degrees, rather than repeatedly evolving the system independently or uniformly losing it in fungi and plants ^23,29^.

In line with this, our homology searches detect SLBP-like proteins not only across eukaryotes but also in a small number of archaeal and bacterial proteins. These distant homologs suggest that the SLBP-type RNA-binding domain represents an ancient RNA-binding module that predates the specialisation of histone mRNA processing in eukaryotes. At the same time, the apparent absence of SLBP or other 3’-end processing factors in some taxa should be interpreted with caution as low quality assemblies, annotation gaps or incomplete sampling may have obscured true homologs.

Interpretation of the fungal patterns necessarily depends on the branching order among early diverging lineages. Some phylogenomic studies include Microsporidia near the base of the fungal tree, or leave their position unresolved^101^. Under this basal placement, the presence in this group of distant SL-like structures together with strong PAS motifs suggests that Microsporidia has been shifted greatly away from SLBP-mediated processing. Our data reinforce the view that the phylogenetic placement of Microsporidia requires further scrutiny. Similarly, recent single-cell genomic phylogenomic analyses place Blastocladiomycota earlier than, or as a sister to, Chytridiomycota^102^. Consistent with this finding, our results show that Blastocladiomycota retain a higher proportion of loci with SL motifs and SLBP orthologs than Chytridiomycota, suggesting that dependence on SL-associated processing decreased along the chytrid and later-diverging fungal lineages.

Identifying SL-like structures outside metazoans is challenging as covariance models trained on canonical metazoan histone SLs perform well within animals but have limited sensitivity for more variable hairpins in fungi, plants and protists. To address this, we implemented a custom detection pipeline combining relaxed sequence motif searches, secondary structure prediction and folding-stability criteria to find candidate SLs and measure their distance from the stop codon. Although this approach increases sensitivity, it remains biased towards structures resembling metazoan SLs. More highly divergent or non-canonical SL structures, such as different loop lengths, may therefore not be detected. Consequently, some clades reported here as lacking histone stem loops may represent false negatives.

A further limitation is that a substantial proportion of histone loci lacked sufficient detected SLBP-SL and/or PAS features for assignment to one of the four defined processing states and were therefore excluded from the state analysis. Accordingly, these comparisons describe trends among classifiable loci rather than the complete complement of histone genes within each lineage.

Finally, our analyses depend on publicly available genome assemblies and predicted gene models, some of which are incomplete. Apparent absence of SLBP, U7-associated factors or ncPAPs in poorly assembled genomes may therefore reflect missing sequence rather than true evolutionary loss. Direct experimental mapping of histone mRNA 3′ ends is not currently available for most of the non-metazoan species examined here, so the precise cleavage sites and mechanisms of polyadenylation remain unresolved. In particular, the polymerase involved and any direct role for SLBP will require experimental testing in tractable early-diverging fungi.

Despite these limitations, our work substantially broadens the evolutionary landscape of histone mRNA 3’-end processing. We show that SLBP proteins, SL structures, U7-snRNP components, noncanonical poly(A) polymerases and PAS-dependent cleavage signals show distinct patterns of retention and loss across eukaryotic lineages. In fungi, these patterns reveal intermediate states between canonical SLBP-SL-U7-dependent processing and PAS-directed cleavage and polyadenylation. More broadly, our findings suggest that specialized RNA-processing systems might evolve through prolonged intermediate states in which RNA-recognition elements persist after the machinery originally coupled to them has begun to disappear. Histone mRNA 3′-end processing therefore provides a model for how RNA-processing systems can be remodelled through progressive decoupling and reassortment of their constituent sequence elements and protein factors.

## Methods

### Identification of putative Stem-Loop Binding Protein orthologs in eukaryotes

To characterize the phylogenetic distribution of SLBPs across eukaryotes, we first identified putative SLBP orthologs in the NCBI non-redundant (NR) protein database using BLASTP^34^ with an E-value threshold of 1e-10 and SLBP sequences reported in *Lopez and Samuelsson* (2008) as queries (**full lists available in the GitHub repository; see Data Availability**). Duplicate sequences were removed post-search. We next used an iterative profile-based HMM search tool, *jackhmmer*^35^. Each full-length SLBP sequence was used as a query against the PDB database (916,370 sequence records; accessed in August 2024). Duplicate sequences were again removed post-search.

We downloaded the corresponding GenBank records for 51,458 hits (51,458 unique genes from 4,252 species) and characterized the presence or absence of the histone RNA hairpin-binding protein RNA-binding domain (RBD; Pfam accession PF15247) by running *rpsblast*^36,37^ against the Conserved Domain database (CDD^38^; accessed May 2024) annotating the RBD as present if the domain hit had an E-value < 1. Further, we searched for conserved RNA-binding domains by HMM-based motif scanning using HMMER (hmmscan^39^ version 3.2.1; against Pfam, implemented with the hmmscan.pl wrapper (http://people.cs.bris.ac.uk/~gough/software/).

By this approach, we identified 5,125 SLBP homologs containing the RBD from 2,329 species. Additionally, 46,305 sequences were initially retrieved as SLBP homologs but lacked PF15247, instead containing other domains, with 40,831 proteins with Ankyrin repeat (ANKYR) domains and 4,901 proteins containing UPA domains. Finally, 28 putative SLBP homologs had no detectable conserved domains at the thresholds used (E-value < 1).

We supplemented this dataset with annotated SLBP homologs from OrthoDB v11^40^ (www.orthodb.org) and with proteins exhibiting the same domain architecture as the histone RNA hairpin-binding protein (**Supplementary Table 1**) from InterPro^41^. This approach identified 219 additional species containing 4,667 SLBP homologs. Species were assigned to higher-order eukaryotic clades according to the NCBI Taxonomy Database. Clade names and species-to-clade mappings are provided in **Supplementary Table S2**.

To distinguish bona fide SLBP orthologs from RBD-positive paralogs, we performed explicit gene-species tree reconciliation. We used seqkit^42^ to remove redundancy, leaving 5,926 RBD-positive proteins from the combined methods above, and confirmed RBD presence by scanning with the Pfam HMM model (PF15247) using hmmscan^35^. The predicted RBD was extracted, aligned with MAFFT^43^, trimmed using TrimAl^44^, and used to infer an approximately maximum-likelihood gene tree (**Supplementary Figure S1**) with IQ-TREE^45^. A corresponding eukaryotic species tree was adopted from the NCBI Taxonomy Database which is broadly consistent with a published eukaryotic tree backbone^46^. After reconciling the SLBP gene tree with the species tree using Treerecs^47^, we identified 2,034 speciation and 3,891 duplication nodes across the reconciled tree. These nodes define the ancestry of 5,926 analyzed sequences. Treerecs further inferred 4,556 orthology relationships and 1,369 paralogy relationships among all RBD-containing proteins.

For species in which SLBP was absent, we verified putative absence using TBLASTN^48^ searches against the NCBI Whole Genome Shotgun (WGS) database. All hits with E-value < 0.05 were downloaded, duplicates removed, and each sequence re-evaluated for RBD presence using *rpsblast*. This additional search step identified 420 more species containing 1,440 SLBP-like sequences, completing our eukaryotic SLBP dataset (**Supplementary Figure S2**).

For each SLBP ortholog, we then assessed conservation of SLIP1-interacting motifs required for translation activation of RD histone mRNAs. The SLIP1-binding segment from *Xenopus laevis* SLBP1, where this interaction has been functionally and structurally characterized, was used as reference^49^. For every SLBP protein, we extracted the N-terminal 150 amino acids and performed local alignment of the target region against the reference segment using Biopython *pairwise2*^50^. We annotated a SLIP1-interacting motif as present if the alignment score was >20 and amino acid identity was >40%. For all motif-positive sequences, we extracted the aligned SLBP segment for downstream multiple sequence alignment and motif logo analysis (**Supplementary Figure S3**).

### Identification of putative SLBP orthologs in prokaryotes

Using the same sequence homology searches and structural domain profiling described above, we searched publicly available prokaryotic datasets for SLBP homologs. BLASTP searches identified 141 candidates in Archaea and 4,210 in Bacteria after removal of duplicates. Domain analyses revealed that only 1 archaeal sequence (Accession: RYG41222.1; *unclassified Archaea*) and 1 bacterial sequence (Accession: OGV26409.1; *Legionellales bacterium*) contained a detectable RBD, respectively.

To ensure comprehensive coverage, we further supplemented these results using sequences retrieved from UniParc, as the corresponding InterPro-curated datasets were no longer available. This additional screening identified three candidates with detectable RBD, including 1 archaeal (*Methanobacteriota archaeon*; UniParc accession: UPI000C600A47) and 2 bacterial candidates (*Crocinitomicaceae bacterium* and *Opitutia bacterium*; UniParc accessions: UPI000C6245EF and UPI000C5FD937).

### Detailed homology survey of SLBPs in fungi

Fungal proteomes and genomes were retrieved from the NCBI Assembly database (accessed 11/02/2025). A curated list of fungal species was used to query NCBI, prioritizing reference and representative assemblies. If no proteome was available, the highest-quality genome assembly was selected based on completeness and assembly level. All sequences were downloaded, organized and validated, resulting in a dataset of 426 proteomes and 94 genome assemblies. For species with genome data only, protein-coding genes were predicted using BRAKER2^51^, generating proteomes for downstream analyses. A Pfam HMM for the SLBP domain (Histone RNA hairpin-binding protein domain, PF15247) was searched against this database using *hmmsearch* (HMMER v3.5, http://hmmer.org/) using the --cut_ga option. Combined with the approach above, we identified a total of 124 fungal species with SLBP hits.

### Identification of canonical histone related protein orthologs (LSM10, LSM11, FLASH, eIF4G, and SLIP1)

We identified LSM10, LSM11, FLASH, eIF4G, and SLIP1 orthologs^10,52–56^ following the same strategy described for SLBP and using sequences reported in *Lopez and Samuelsson* (2008) as queries (**full lists available in the GitHub repository; see Data Availability**). Searches were conducted using BLASTP and TBLASTN with E-value < 1e-5, and functional domains were tested using both *rpsblast* and *hmmscan*. To expand the dataset, annotated sequences from external databases were included (OrthoDB v12.1, CDD, and InterPro; **Supplementary Table S1**).

For FLASH, additional domain-specific filtering was required because the relevant functional domain (CDD: cd12202) is shared among unrelated proteins^57^. We used the PANTHER HMM profile PTHR15489 to identify FLASH candidates via *hmmscan*^58^, and further tested all candidates against the PANTHER model PTHR16088 (YY1-associated transcriptional regulators). To identify bona fide FLASH orthologs, we first applied both HMM profiles to all sequences and then set a threshold of E-value < 1e-5 to filter out FLASH candidates. Among these, sequences were removed as likely GON-4L homologs if their match to PTHR16088 were at least 1,000-fold more significant than their match to PTHR15489.

To identify SLIP1 sequences, we did not use simple domain annotation, as the MIF4G domain is shared by multiple unrelated proteins (e.g. eIF4G, eIF-2B, UPF3, and nuclear cap-binding proteins like CBP80, CBC1 and NCBP1)^59–63^. Instead, we implemented a multi-step filtering and phylogenetic validation approach. First, we extracted all proteins containing the MIF4G domain, and filtered for complete sequences shorter than 300 amino acids, excluding longer eIF4G-like isoforms and CTIF (cap binding complex dependent translation initiation factor). Next, candidates were compared against PTHR23253 (Eukaryotic Translation Initiation Factor 4 Gamma) and PTHR23254 (mRNA Translation Regulation / Replication-Dependent Histone mRNA Translation) HMM profiles. After applying a baseline significance threshold of E-value < 1e-5, sequences with at least 1,000-fold more significant match to PTHR23254 were retained as candidate SLIP1 homologs. We further examined the motifs in SLIP1 implicated in SLBP interaction^64^. Retained sequences were aligned with MAFFT ^43^ using default parameters, and an approximately maximum-likelihood phylogeny (**Supplementary Figure S4)** was inferred using FastTree^65^. Sequences clustering with known SLIP1 references in the resulting tree were classified as SLIP1-like proteins.

### Survey of histone proteins in fungi, plants, protists, and metazoans

To characterize histone proteins across major eukaryotic clades, we retrieved protein-coding sequences (CDS) from the NCBI protein database using the keyword “histone” (accessed November 2024). For all retrieved CDS, we identified putative histone genes by running *rpstblastn*^37^ to search for histone-fold domains (HFDs) ^66–69^ against CDD (full list in **Supplementary Table S3**), retaining hits with E-value < 1e-5. To check the completeness of the coding sequence, we first filtered sequences based on NCBI metadata annotations. We then examined each sequence for the presence of a canonical start codon (ATG) and one of the standard stop codons (TAA, TAG and TGA). We excluded sequences with loss of the 3’-end stop codon or loss of both 3’ and 5’ ends, as no downstream genomic region could be retrieved relative to the annotated stop codon. We excluded low-confidence annotations (e.g. sequences derived from metagenome-assembled genomes (MAGs) or annotated as linker histones or “histone-replacement-like”. Full list of exclusions provided in **Supplementary Table S4**).

#### Metazoans

We retrieved 400,271 metazoan CDS, of which 201,398 contained a detectable HFD. We excluded 75,042 incomplete sequences (lacking both stop and start codons), as well as 5,419 sequences lacking 3’ stop codons only. We removed an additional 293 low-confidence entries, resulting in 101,518 high-confidence metazoan CDS, among which 69,876 had at least 26 bp retrievable downstream flanking sequences. This 26 bp threshold ensures the inclusion of the entire stem loop structure, which comprises a 5 nt space before the stem loop, a 16 nt stem loop and a 4-5 bp downstream space for cleavage^29^ (per-sequence metadata in **Supplementary File 1**).

#### Fungi

We retrieved 154,626 fungal CDS, among which 54,267 contained a detectable HFD. We removed 20,904 incomplete sequences and 415 sequences lacking a 3’ stop codon. We filtered out an additional 640 low-confidence sequences, resulting in 32,303 high-confidence fungal histone CDS (**Supplementary File 2**).

#### Plants

We retrieved 67,399 CDS, with 28,010 containing an HFD. We excluded 795 incomplete sequences and 240 sequences lacking a 3’ stop codon. We further removed 977 low-confidence sequences, resulting in a final 25,998 plant histone CDS (**Supplementary File 3**).

#### Protists

We retrieved 17,535 CDS sequences for all protists for 17,535 entries, with 16,809 containing HFD domains. We initially removed 1,360 low confidence sequences. After restricting to protozoan species (Alveolata, Stramenopiles, Rhizaria, Amoebozoa, Metamonada, Hemimastigophora, Discoba, and Ichthyosporea), 8,068 sequences remained. We excluded 288 incomplete sequences and 345 sequences missing a 3’ stop codon, resulting in a final dataset of 7,435 histone CDS (**Supplementary File 4**).

### Identification of histone mRNA 3’ UTRs stem-loops and U7 snRNA within metazoa

To detect histone stem-loops, we used the Rfam covariance model (RF00032) and ran *cmsearch*^70^ against the 300 bp flanking sequences downstream of histone mRNA. This approach accounts for both sequence conservation and RNA secondary structure to detect canonical histone SL motifs. As SL structures in metazoans are highly conserved and well-curated in Rfam, covariance model searches provide a robust and reliable strategy for identifying SL candidates in this group.

For U7 snRNA, we retrieved annotated sequences from Rfam (RF00066), which includes curated U7 snRNA entries across a wide range of metazoan species. Because replication-dependent histone mRNA processing and the associated SL and U7 elements are already well established in metazoans (see Marzluff et al. 2008), we did not perform exhaustive genome-wide detection in this group. Instead, curated Rfam annotations were used to establish a reference for comparison with other eukaryotic clades. This approach also reflects practical considerations, as comprehensive genome-scale screening across all metazoan species was not computationally feasible.

### Detection of histone mRNA 3’ UTRs stem-loops and U7 snRNA outside metazoa

Outside metazoans, histone SL structures are more variable and can diverge substantially from canonical motifs. To identify potentially divergent stem-loop structures in histone mRNA 3’ UTRs, we used a two-step approach. First, we scanned for sequences consistent with a canonical stem-loop structure (6 bp stem, 4 nucleotide loop). Second, we used RNAfold^71,72^ to confirm the secondary structure.

We used *efetch*^73^ from the NCBI toolkit to retrieve 300 bp downstream of histone stop codons. Each sequence was scanned using a sliding window of 16 nucleotides that included a 6 base 5’ stem, a 4 nucleotide loop, and 6 base 3’ stem^74^. We enforced Watson-Crick pairing rules and allowed one mismatch for all possible candidates. If a candidate SL was detected, the sequence was extracted with 5 additional nucleotides on each side (26 nucleotides in total) for subsequent secondary structure confirmation.

RNAfold was then applied to predict the secondary structure of each 26 nt sequence^71,72^. Only sequences that folded into the expected SL structure (i.e. “(((((())))))” in dot-bracket notation) were retained. When multiple candidate SLs were detected within a histone downstream region, the structure with the lowest predicted minimum free energy was retained.

To assess the reproducibility of this approach, we tested our custom script against the curated SLs in Rfam (RF00032). Our method recovered over 99% of SL entries (44,610 / 44,646). The 36 missed SLs corresponded to non-canonical SL structures with 3- or 5-nt loops, two or more mismatches in stem, or sequences that failed to form a recognizable SL structure according to RNAfold (**Supplementary Table S5**). To identify purine-rich HDEs^19^, we scanned for recurring motifs (“AAA” or “AAGA”) in downstream sequences. Using this approach, purine-rich regions present in all RF00032 sequences from Rfam could be successfully recovered.

To detect putative U7 snRNA across fungal, plant, and protist genomes, we downloaded 9,537 reference genomes (one reference per species) from NCBI Assembly, comprising 5,352 fungal, 3,424 plant, and 761 protist genomes. We used the Rfam covariance model (RF00066) and ran *cmsearch*^70^ against these genomes. The search was conducted with a reporting bit score threshold of 23.34, as specified for the U7 model in the Rfam database.

### Correlation of Sequence Conservation and Distance Analysis

For each clade, we quantified how SL conservation varied with distance from the histone stop codon. Histone entries were excluded if the SL is missing, not exactly 26 bp in length, or containing ambiguous bases (for example, “N” and “Y”) The 16-nt stem–loop motif corresponding to the predicted hairpin (6-bp stem, 4-nt loop, 6-bp stem) was then extracted. Conservation was scored as exact matches to the reference (from the *hmmscan* best hit: XP_010200268.2, representing histone H2A from *Colius striatus*). Extreme distances were excluded by restricting distance to 20-200 nt. Distances were binned in 10-nt intervals and mean conservation was computed per bin. Associations between distance and conservation were assessed on binned means using Pearson’s r and Spearman’s rho implemented in SciPy^75^.

### Stem-Loop Similarity Network Construction

To summarise sequence space, we extracted the central 16-nt cores from each 26-nt SL. Identical cores were grouped together as unique variants, and their frequencies were counted. We then constructed undirected similarity graphs in NetworkX^76^. Nodes in the network represent unique cores. Edges connect pairs of cores based on loop edit distance, defined as the number of nucleotide changes required to convert one sequence to another. To highlight the most frequent SL variants, we built a top 50 subgraph using the 50 most frequent cores. For display, node labels included predicted folding free energies (delta G, kcal/mol; **Supplementary Figure S7A-D**) computed with NUPACK^77^ in Python under the RNA model at 27°C. Different colours were used to indicate exclusivity (e.g. Bilateria-specific vs variants shared across other metazoan groups). All preprocessing, binning, correlation, and network parameters were held constant across metazoans, Protozoa, fungi and plants.

### Quantification of SLBP-SL/PAS processing modes in basal fungi

To compare histone 3’-end processing modes across early-diverging fungal clades, we scanned for canonical poly(A) signal motif (AATAAA; RNA equivalent: AAUAAA) and its variants (ATTAAA, AGTAAA, TATAAA, CATAAA, GATAAA, AATACA, AATATA and AATAGA) ^78–80^ across the 300-nt downstream flanking sequences. A locus was called PAS-positive if a PAS signal was present in the 300 bp downstream sequence. Each locus was assigned to one of four mutually exclusive states: SLBP-SL, SLBP-SL-PAS, SLBP-PAS or PAS-only. Protein records not meeting any of these definitions were excluded from state analysis (35,869/52,444). Thus, the state analysis was restricted to loci containing sufficient detected features to distinguish among the defined SL- and/or PAS-associated states. For each species, we calculated the fraction of loci in each state (summing to 1). For each clade, we reported the means of these species-level fractions, the dominant state per species (largest fraction; ties resolved to the coexistence class SLBP-SL-PAS), and a clade-wise contrast between SL-containing modes and PAS-only: for each species we set p_SL_ = (SLBP-SL +SLBP-SL-PAS) and p_PAS_ = (PAS-only); then estimated logit(p_SL_) - logit(p_PAS_) with 95% confidence intervals by nonparametric bootstrap across species.

### Identification of Candidate GLD2-family Polymerases in Fungi

To identify putative cytoplasmic non-canonical poly(A) polymerases related to GLD2 in fungi, we performed protein similarity searches against NCBI nr (as above), yielding 11,140 protein sequences for downstream analysis. Incomplete entries were removed based on “partial” keyword annotations in FASTA headers. We then used a panel of InterPro HMMs (**Supplementary Table S6**) to screen and remove unrelated nucleotidyltransferases. To enrich non-canonical polymerases, we scanned remaining candidates using PTHR12271(DNA polymerase type-B-like RNA processing) and PTHR23092 (POLY(A) RNA POLYMERASE) and only retained sequences with domain bit-score > 200, leaving 5,685 and 3,382 candidates, respectively.

To distinguish poly(A) from poly(U)-adding polymerases, we classified all candidate proteins using a panel of subfamily-specific HMMs representing GLD2-like polymerases, the major *Schizosaccharomyces pombe* Cid subfamilies (cid1, cid11, cid13) as well as trf4-like enzymes. Previous structural and biochemical studies have shown that nucleotide specificity in GLD2 is influenced by residues surrounding the catalytic site. In particular, a histidine (H) residue located near the GLD2 T439-N440 region (corresponding to Cid1 H336) shifts GLD2 from ATP-selective to UTP-selective^31,32^. We therefore aligned the catalytic core regions of all candidate sequences to the human GLD2 reference and examined a ±2 residue window surrounding the corresponding position. Proteins containing histidine at this position were classified as likely U-adding polymerases, whereas those lacking histidine were classified as candidate A-adding ncPAPs.

### Structural modelling of GLD2-like proteins

For structural comparison, a representative fungal GLD2-like candidate (NCBI: KAI8803508.1) and human GLD-2 (UniProt: Q6PIY7) were modelled with AlphaFold2-ptm ^81,82^. Per-residue model confidence was inferred from pLDDT scores encoded in the predicted structures.

Structural alignment was performed in UCSF ChimeraX^83^ using the MatchMaker algorithm with default parameters. Structural conservation of the nucleotidyltransferase catalytic region and the ATP/UTP specificity-determining region surrounding the human GLD2 T439-N440 segment was examined in the resulting superposition. For visualization, extended low-confidence terminal regions of the fungal GLD2-like model were trimmed to improve the clarity of the structure.

### Survey of histone transcript termini and expression support in fungal RNA-seq datasets

To investigate histone mRNA 3’ end formation in fungi, we surveyed transcript termini across seven RNA-seq datasets (**Supplementary Table S7**). Our primary objective was to identify SLBP-mediated cleavage sites. We retrieved raw sequences from the NCBI Sequence Read Archive using *prefetch*^84^. Reads were then aligned to the corresponding reference genomes using HISAT2^85^, and resulting alignments were converted to sorted BAM files using SAMtools^86^. Alignments were visualized in Integrative Genome Viewer (IGV)^87^ to assess reads coverage across histone CDS and downstream regions.

For each RNA-seq dataset, we screened for poly(A) tails to further assess polyadenylation. FASTQ files were searched for sequences matching histone CDS and downstream flanking sequences. We used direct string matching to identify reads containing histone-specific sequence followed by candidate poly(A) tails. Only poly(A) tracts of at least 15 nt were retained to avoid potential sequencing artefacts^88^.

To test whether fungal histone processing components were expressed in the same developmental context, we additionally analysed four RNA-seq datasets (**Supplementary Table S8**) from *Spizellomyces punctatus*. Raw reads were aligned to the *S. punctatus* reference genome (NCBI: GCF_000182565.1) using HISAT2. The protein sequence of the ncPAP candidate (NCBI: XP_016605801.1) was mapped back to the reference genome using TBLASTN. Seven core histones CDS and SLBP CDS sequences (**Supplementary Table S9**) were mapped to the *S. punctatus* reference genome using BLASTN. The resulting coding intervals were converted to BED format. Then only the primary mapped reads overlapping with corresponding CDS intervals were counted from the HISAT2 alignments using SAMtools. Furthermore, read counts were normalized as counts per million primary mapped reads (CPM). Finally, fold changes were calculated between zoospore and multinucleated groups for candidate ncPAP, core histones and SLBP.

## Author Contributions

M.N.L. and U.B. conceived the project. B.Z. analysed data and produced all main and supplemental figures. M.N.L. and B.Z. interpreted data and wrote the manuscript. M.N.L. supervised the project and assisted with data interpretation.

## Acknowledgements

We thank Carsten Kröger for useful comments on the manuscript, and Damien Woods and Ahmed Shalaby for helpful discussions and advice regarding NUPACK-based RNA folding predictions and the evaluation of stem-loop stability.

## Data Availability

Code generated for this study can be found in GitHub at https://github.com/BZhao95/SLBP_Histone_Evolution. All other study data are included in the article and/or supporting information.

## Supplementary Figures

**Supplementary Figure S1:** Phylogenetic tree of the SLBP RNA-binding domain.

**Supplementary Figure S2:** Public sources of SLBP ortholog annotation across eukaryotes.

**Supplementary Figure S3:** Sequence conservation of the SLBP SLIP1-interacting motif.

**Supplementary Figure S4:** Phylogeny of SLIP1-like MIF4G-domain proteins.

**Supplementary Figure S5:** Sequence similarity between *Takifugu rubripes* FLASH-associated domain query and translated genomic region in *Rozella allomycis*.

**Supplementary Figure S6**: Identification and secondary-structure validation of a U7-like snRNA in *Rozella allomycis*.

**Supplementary Figure S7A:** Similarity network of the 50 most frequent metazoan histone stem loops.

**Supplementary Figure S7B:** Similarity network of the 50 most frequent fungal histone stem loops.

**Supplementary Figure S7C:** Similarity network of the 50 most frequent protist histone stem loops.

**Supplementary Figure S7D:** Similarity network of the 50 most frequent plant histone stem loops.

**Supplementary Figure S8:** Public mRNA-seq datasets support predicted locus-level 3′-end processing patterns.

**Supplementary Figure S9:** Expression of candidate SLBP-associated processing factors and core histones in *Spizellomyces punctatus*.

## Supplementary Tables

**Supplementary Table S1:** External database annotations and HMM profiles used for ortholog identification.

**Supplementary Table S2:** NCBI Taxonomy clades with and without SLBP.

**Supplementary Table S3:** NCBI Conserved Domain Database (CDD) entries used for identification of canonical core histone proteins.

**Supplementary Table S4:** NCBI Annotation terms used to exclude non-canonical histone-domain-containing proteins.

**Supplementary Table S5:** Rfam-curated histone stem-loop sequences not recovered by the custom script.

**Supplementary Table S6:** Domain profiles used to filter unrelated nucleotidyltransferase and RNA-binding proteins.

**Supplementary Table S7:** Fungal RNA-seq datasets surveyed for histone transcript termini.

**Supplementary Table S8:** RNA-seq datasets used to assess expression of histone 3’ end-processing factors and core histone genes in Spizellomyces punctatus DAOM BR117.

**Supplementary Table S9:** Candidate processing factors and histone genes used for expression analysis.

**Supplementary Table S10:** Median histone stem-loop distances and mean conservation scores across major clades.

## Supplementary Files

**Supplementary File 1. Metazoan histone CDS survey dataset.** Survey of metazoan histone genes and associated 3’-end processing features. Data include protein accessions, conserved domain annotations, species and taxonomic information, presence or absence of SLBP, histone protein annotations, detection of stem-loop motifs, SL position relative to the stop codon, and predicted SL sequences.

**Supplementary File 2. Fungi histone CDS survey dataset.** Survey of fungal histone genes and associated 3’-end processing features. Data include protein accessions, conserved domain annotations, species and taxonomic information, presence or absence of SLBP, histone protein annotations, detection of stem-loop motifs, SL position relative to the stop codon, and predicted SL sequences.

**Supplementary File 3. Plant histone CDS survey dataset.** Survey of plant histone genes and associated 3’-end processing features. Data include protein accessions, conserved domain annotations, species and taxonomic information, presence or absence of SLBP, histone protein annotations, detection of stem-loop motifs, SL position relative to the stop codon, and predicted SL sequences.

**Supplementary File 4. Protozoan histone CDS survey dataset.** Survey of protozoan histone genes and associated 3’-end processing features. Data include protein accessions, conserved domain annotations, species and taxonomic information, presence or absence of SLBP, histone protein annotations, detection of stem-loop motifs, SL position relative to the stop codon, and predicted SL sequences.

## References

1. Marzluff, W. F. Histone 3’ ends: essential and regulatory functions. Gene Expr. 2, 93–97 (1992).

2. Davin Townley-Tilson, W. H., Pendergrass, S. A., Marzluff, W. F. & Whitfield, M. L. Genome-wide analysis of mRNAs bound to the histone stem–loop binding protein. RNA 12, 1853–1867 (2006).

3. Zanier, K. et al. Structure of the histone mRNA hairpin required for cell cycle regulation of histone gene expression. RNA 8, 29–46 (2002).

4. Zhang, M., Lam, T. T., Tonelli, M., Marzluff, W. F. & Thapar, R. Interaction of the histone mRNA hairpin with stem-loop binding protein (SLBP) and regulation of the SLBP-RNA complex by phosphorylation and proline isomerization. Biochemistry 51, 3215–3231 (2012).

5. Sullivan, K. D., Mullen, T. E., Marzluff, W. F. & Wagner, E. J. Knockdown of SLBP results in nuclear retention of histone mRNA. RNA 15, 459–472 (2009).

6. Whitfield, M. L. et al. Stem-loop binding protein, the protein that binds the 3’ end of histone mRNA, is cell cycle regulated by both translational and posttranslational mechanisms. Mol Cell Biol 20, 4188–4198 (2000).

7. Wang, Z. F., Whitfield, M. L., Ingledue, T. C., 3rd, Dominski, Z. & Marzluff, W. F. The protein that binds the 3’ end of histone mRNA: a novel RNA-binding protein required for histone pre-mRNA processing. Genes Dev. 10, 3028–3040 (1996).

8. Martin, F., Schaller, A., Eglite, S., Schümperli, D. & Müller, B. The gene for histone RNA hairpin binding protein is located on human chromosome 4 and encodes a novel type of RNA binding protein. EMBO J. 16, 769–778 (1997).

9. Dominski, Z., Yang, X.-C. & Marzluff, W. F. The polyadenylation factor CPSF-73 is involved in histone-pre-mRNA processing. Cell 123, 37–48 (2005).

10. Yang, X.-C., Burch, B. D., Yan, Y., Marzluff, W. F. & Dominski, Z. FLASH, a proapoptotic protein involved in activation of caspase-8, is essential for 3’ end processing of histone pre-mRNAs. Mol. Cell 36, 267–278 (2009).

11. Kolev, N. G. & Steitz, J. A. Symplekin and multiple other polyadenylation factors participate in 3’-end maturation of histone mRNAs. Genes Dev 19, 2583–2592 (2005).

12. Barcaroli, D. et al. FLASH is required for histone transcription and S-phase progression. Proc. Natl. Acad. Sci. U. S. A. 103, 14808–14812 (2006).

13. Xiang, K. et al. Crystal structure of the human symplekin-Ssu72-CTD phosphopeptide complex. Nature 467, 729–733 (2010).

14. Kolev, N. G., Yario, T. A., Benson, E. & Steitz, J. A. Conserved motifs in both CPSF73 and CPSF100 are required to assemble the active endonuclease for histone mRNA 3’-end maturation. EMBO Rep. 9, 1013–1018 (2008).

15. Ryan, K., Calvo, O. & Manley, J. L. Evidence that polyadenylation factor CPSF-73 is the mRNA 3’ processing endonuclease. RNA 10, 565–573 (2004).

16. Kaufmann, I., Martin, G., Friedlein, A., Langen, H. & Keller, W. Human Fip1 is a subunit of CPSF that binds to U-rich RNA elements and stimulates poly(A) polymerase. EMBO J. 23, 616–626 (2004).

17. Yao, Y., Song, L., Katz, Y. & Galili, G. Cloning and characterization of Arabidopsis homologues of the animal CstF complex that regulates 3’ mRNA cleavage and polyadenylation. J. Exp. Bot. 53, 2277–2278 (2002).

18. Spycher, C. et al. 3′ End processing of mouse histone pre-mRNA: evidence for additional base-pairing between U7 snRNA and pre-mRNA. Nucleic Acids Res. 22, 4023–4030 (1994).

19. Bond, U. M., Yario, T. A. & Steitz, J. A. Multiple processing-defective mutations in a mammalian histone pre-mRNA are suppressed by compensatory changes in U7 RNA both in vivo and in vitro. Genes Dev 5, 1709–1722 (1991).

20. Marzluff, W. F. & Duronio, R. J. Histone mRNA expression: multiple levels of cell cycle regulation and important developmental consequences. Curr. Opin. Cell Biol. 14, 692–699 (2002).

21. Dominski, Z. et al. 3’ end processing of Drosophila melanogaster histone pre-mRNAs: requirement for phosphorylated Drosophila stem-loop binding protein and coevolution of the histone pre-mRNA processing system. Mol. Cell. Biol. 22, 6648–6660 (2002).

22. Wagner, E. J. et al. Conserved zinc fingers mediate multiple functions of ZFP100, a U7snRNP associated protein. RNA 12, 1206–1218 (2006).

23. López, M. D. & Samuelsson, T. Early evolution of histone mRNA 3′ end processing. RNA 14, 1–10 (2008).

24. Marzluff, W. F. Metazoan replication-dependent histone mRNAs: a distinct set of RNA polymerase II transcripts. Curr. Opin. Cell Biol. 17, 274–280 (2005).

25. Dominski, Z. & Marzluff, W. F. Formation of the 3′ end of histone mRNA: Getting closer to the end. Gene 396, 373–390 (2007).

26. He, S. et al. SUMO2 promotes histone pre-mRNA processing by stabilizing histone locus body interactions and facilitating U7 snRNP assembly. Genes Dev 39, 1182–1197 (2025).

27. Pettinati, I. et al. Biosynthesis of histone messenger RNA employs a specific 3’ end endonuclease. Elife 7, (2018).

28. Marzluff, W. F. & Koreski, K. P. Birth and death of histone mRNAs. Trends Genet. 33, 745–759 (2017).

29. Marzluff, W. F., Wagner, E. J. & Duronio, R. J. Metabolism and regulation of canonical histone mRNAs: life without a poly(A) tail. Nat. Rev. Genet. 9, 843–854 (2008).

30. Battle, D. J. & Doudna, J. A. The stem-loop binding protein forms a highly stable and specific complex with the 3′ stem-loop of histone mRNAs. RNA 7, 123–132 (2001).

31. Nakel, K., Bonneau, F., Eckmann, C. R. & Conti, E. Structural basis for the activation of the C. elegans noncanonical cytoplasmic poly(A)-polymerase GLD-2 by GLD-3. Proc. Natl. Acad. Sci. U. S. A. 112, 8614–8619 (2015).

32. Chung, C. Z., Jo, D. H. S. & Heinemann, I. U. Nucleotide specificity of the human terminal nucleotidyltransferase Gld2 (TUT2). RNA 22, 1239–1249 (2016).

33. Pérez-Roldán, J. et al. Maternal histone mRNAs are uniquely processed through polyadenylation in a Stem-Loop Binding Protein (SLBP) dependent manner. Nucleic Acids Res. 53, gkaf288 (2025).

34. Altschul, S. F., Gish, W., Miller, W., Myers, E. W. & Lipman, D. J. Basic local alignment search tool. J Mol Biol 215, 403–410 (1990).

35. Eddy, S. R. A new generation of homology search tools based on probabilistic inference. Genome Inform. 23, 205–211 (2009).

36. Boratyn, G. M. et al. Domain enhanced lookup time accelerated BLAST. Biol. Direct 7, 12 (2012).

37. Camacho, C. et al. BLAST+: architecture and applications. BMC Bioinformatics 10, 421 (2009).

38. Marchler-Bauer, A. et al. CDD: NCBI’s conserved domain database. Nucleic Acids Res. 43, D222–D226 (2014).

39. Finn, R. D., Clements, J. & Eddy, S. R. HMMER web server: interactive sequence similarity searching. Nucleic Acids Res. 39, W29–37 (2011).

40. Kuznetsov, D. et al. OrthoDB v11: annotation of orthologs in the widest sampling of organismal diversity. Nucleic Acids Res. 51, D445–D451 (2023).

41. Paysan-Lafosse, T. et al. InterPro in 2022. Nucleic Acids Res. 51, D418–D427 (2023).

42. Shen, W., Le, S., Li, Y. & Hu, F. SeqKit: A cross-platform and ultrafast toolkit for FASTA/Q file manipulation. PLoS One 11, e0163962 (2016).

43. Katoh, K. & Standley, D. M. MAFFT multiple sequence alignment software version 7: improvements in performance and usability. Mol Biol Evol 30, 772–780 (2013).

44. Capella-Gutiérrez, S., Silla-Martínez, J. M. & Gabaldón, T. trimAl: a tool for automated alignment trimming in large-scale phylogenetic analyses. Bioinformatics 25, 1972–1973 (2009).

45. Minh, B. Q. et al. IQ-TREE 2: New models and efficient methods for phylogenetic inference in the genomic era. Mol. Biol. Evol. 37, 1530–1534 (2020).

46. Burki, F., Roger, A. J., Brown, M. W. & Simpson, A. G. B. The new tree of eukaryotes. Trends Ecol. Evol. 35, 43–55 (2020).

47. Comte, N. et al. Treerecs: an integrated phylogenetic tool, from sequences to reconciliations. Bioinformatics 36, 4822–4824 (2020).

48. Gertz, E. M., Yu, Y.-K., Agarwala, R., Schäffer, A. A. & Altschul, S. F. Composition-based statistics and translated nucleotide searches: improving the TBLASTN module of BLAST. BMC Biol. 4, 41 (2006).

49. Cakmakci, N. G., Lerner, R. S., Wagner, E. J., Zheng, L. & Marzluff, W. F. SLIP1, a factor required for activation of histone mRNA translation by the stem-loop binding protein. Mol. Cell. Biol. 28, 1182–1194 (2008).

50. Cock, P. J. A. et al. Biopython: freely available Python tools for computational molecular biology and bioinformatics. Bioinformatics 25, 1422–1423 (2009).

51. Brůna, T., Hoff, K. J., Lomsadze, A., Stanke, M. & Borodovsky, M. BRAKER2: automatic eukaryotic genome annotation with GeneMark-EP+ and AUGUSTUS supported by a protein database. NAR Genom. Bioinform. 3, lqaa108 (2021).

52. Wagner, E. J. & Marzluff, W. F. ZFP100, a component of the active U7 snRNP limiting for histone pre-mRNA processing, is required for entry into S phase. Mol Cell Biol 26, 6702–6712 (2006).

53. Pillai, R. S., Will, C. L., Lührmann, R., Schümperli, D. & Müller, B. Purified U7 snRNPs lack the Sm proteins D1 and D2 but contain Lsm10, a new 14 kDa Sm D1-like protein. EMBO J. 20, 5470–5479 (2001).

54. Gross, J. D. et al. Ribosome loading onto the mRNA cap is driven by conformational coupling between eIF4G and eIF4E. Cell 115, 739–750 (2003).

55. Godfrey, A. C., White, A. E., Tatomer, D. C., Marzluff, W. F. & Duronio, R. J. The Drosophila U7 snRNP proteins Lsm10 and Lsm11 are required for histone pre-mRNA processing and play an essential role in development. RNA 15, 1661–1672 (2009).

56. Yoshii, M. et al. The Arabidopsis cucumovirus multiplication 1 and 2 loci encode translation initiation factors 4E and 4G. J. Virol. 78, 6102–6111 (2004).

57. Lu, P. et al. The developmental regulator protein Gon4l associates with protein YY1, co-repressor Sin3a, and histone deacetylase 1 and mediates transcriptional repression. J. Biol. Chem. 286, 18311–18319 (2011).

58. Eddy, S. R. A probabilistic model of local sequence alignment that simplifies statistical significance estimation. PLoS Comput. Biol. 4, e1000069 (2008).

59. Aravind, L. & Koonin, E. V. Eukaryote-specific domains in translation initiation factors: implications for translation regulation and evolution of the translation system. Genome Res 10, 1172–1184 (2000).

60. Worch, R. et al. Specificity of recognition of mRNA 5’ cap by human nuclear cap-binding complex. RNA 11, 1355–1363 (2005).

61. Mazza, C., Ohno, M., Segref, A., Mattaj, I. W. & Cusack, S. Crystal structure of the human nuclear cap binding complex. Mol Cell 8, 383–396 (2001).

62. Gunawardena, S. R., Ruis, B. L., Meyer, J. A., Kapoor, M. & Conklin, K. F. NOM1 targets protein phosphatase I to the nucleolus. J. Biol. Chem. 283, 398–404 (2008).

63. Levy-Strumpf, N., Deiss, L. P., Berissi, H. & Kimchi, A. DAP-5, a novel homolog of eukaryotic translation initiation factor 4G isolated as a putative modulator of gamma interferon-induced programmed cell death. Mol. Cell. Biol. 17, 1615–1625 (1997).

64. von Moeller, H. et al. Structural and biochemical studies of SLIP1-SLBP identify DBP5 and eIF3g as SLIP1-binding proteins. Nucleic Acids Res 41, 7960–7971 (2013).

65. Price, M. N., Dehal, P. S. & Arkin, A. P. FastTree 2--approximately maximum-likelihood trees for large alignments. PLoS One 5, e9490 (2010).

66. Bhasin, M., Reinherz, E. L. & Reche, P. A. Recognition and classification of histones using support vector machine. J. Comput. Biol. 13, 102–112 (2006).

67. Baxevanis, A. D., Arents, G., Moudrianakis, E. N. & Landsman, D. A variety of DNA-binding and multimeric proteins contain the histone fold motif. Nucleic Acids Res. 23, 2685–2691 (1995).

68. Baxevanis, A. D. & Landsman, D. Histone and histone fold sequences and structures: a database. Nucleic Acids Res. 25, 272–273 (1997).

69. Thatcher, T. H. & Gorovsky, M. A. Phylogenetic analysis of the core histones H2A, H2B, H3, and H4. Nucleic Acids Res. 22, 174–179 (1994).

70. Nawrocki, E. P. & Eddy, S. R. Infernal 1.1: 100-fold faster RNA homology searches. Bioinformatics 29, 2933–2935 (2013).

71. Gruber, A. R., Lorenz, R., Bernhart, S. H., Neuböck, R. & Hofacker, I. L. The Vienna RNA websuite. Nucleic Acids Res. 36, W70–4 (2008).

72. Lorenz, R. et al. ViennaRNA Package 2.0. Algorithms Mol. Biol. 6, 26 (2011).

73. Sayers, E. The E-utilities in-depth: parameters, syntax and more. Entrez Programming Utilities Help [Internet] https://www.ncbi.nlm.nih.gov/books/NBK25499/ (2009).

74. Pandey, N. B. & Marzluff, W. F. The stem-loop structure at the 3’ end of histone mRNA is necessary and sufficient for regulation of histone mRNA stability. Mol. Cell. Biol. 7, 4557–4559 (1987).

75. Virtanen, P. et al. SciPy 1.0: fundamental algorithms for scientific computing in Python. Nat. Methods 17, 261–272 (2020).

76. Hagberg, A. A., Schult, D. A. & Swart, P. J. Exploring network structure, dynamics, and function using NetworkX. in Proceedings of the Python in Science Conference 11–15 (SciPy, 2008).

77. Zadeh, J. N. et al. NUPACK: Analysis and design of nucleic acid systems. J. Comput. Chem. 32, 170–173 (2011).

78. van Helden, J., del Olmo, M. & Pérez-Ortín, J. E. Statistical analysis of yeast genomic downstream sequences reveals putative polyadenylation signals. Nucleic Acids Res. 28, 1000–1010 (2000).

79. Graber, J. H., Cantor, C. R., Mohr, S. C. & Smith, T. F. In silico detection of control signals: mRNA 3’-end-processing sequences in diverse species. Proc. Natl. Acad. Sci. U. S. A. 96, 14055–14060 (1999).

80. Stroup, E. K. & Ji, Z. Delineating yeast cleavage and polyadenylation signals using deep learning. Genome Res. 34, 1066–1080 (2024).

81. Jumper, J. et al. Highly accurate protein structure prediction with AlphaFold. Nature 596, 583–589 (2021).

82. Varadi, M. et al. AlphaFold Protein Structure Database in 2024: providing structure coverage for over 214 million protein sequences. Nucleic Acids Res. 52, D368–D375 (2024).

83. Pettersen, E. F. et al. UCSF ChimeraX: Structure visualization for researchers, educators, and developers. Protein Sci. 30, 70–82 (2021).

84. Leinonen, R., Sugawara, H., Shumway, M. & International Nucleotide Sequence Database Collaboration. The sequence read archive. Nucleic Acids Res. 39, D19–21 (2011).

85. Kim, D., Paggi, J. M., Park, C., Bennett, C. & Salzberg, S. L. Graph-based genome alignment and genotyping with HISAT2 and HISAT-genotype. Nat. Biotechnol. 37, 907–915 (2019).

86. Li, H. et al. The Sequence Alignment/Map format and SAMtools. Bioinformatics 25, 2078–2079 (2009).

87. Robinson, J. T. et al. Integrative genomics viewer. Nat. Biotechnol. 29, 24–26 (2011).

88. Svoboda, M., Frost, H. R. & Bosco, G. Internal oligo(dT) priming introduces systematic bias in bulk and single-cell RNA sequencing count data. NAR Genom. Bioinform. 4, lqac035 (2022).

89. Hocher, A. et al. Histones with an unconventional DNA-binding mode in vitro are major chromatin constituents in the bacterium Bdellovibrio bacteriovorus. Nat. Microbiol. 8, 2006–2019 (2023).

90. Hu, Y. et al. Bacterial histone HBb from Bdellovibrio bacteriovorus compacts DNA by bending. Nucleic Acids Res. 52, 8193–8204 (2024).

91. Irwin, N. A. T. & Richards, T. A. Self-assembling viral histones are evolutionary intermediates between archaeal and eukaryotic nucleosomes. Nat. Microbiol. 9, 1713–1724 (2024).

92. Liu, Y. et al. Virus-encoded histone doublets are essential and form nucleosome-like structures. Cell 184, 4237–4250.e19 (2021).

93. Letunic, I. & Bork, P. Interactive Tree Of Life (iTOL) v5: an online tool for phylogenetic tree display and annotation. Nucleic Acids Res. 49, W293–W296 (2021).

94. Beggs, S., James, T. C. & Bond, U. The PolyA tail length of yeast histone mRNAs varies during the cell cycle and is influenced by Sen1p and Rrp6p. Nucleic Acids Res. 40, 2700–2711 (2012).

95. Reis, C. C. & Campbell, J. L. Contribution of Trf4/5 and the nuclear exosome to genome stability through regulation of histone mRNA levels in Saccharomyces cerevisiae. Genetics 175, 993–1010 (2007).

96. Canavan, R. & Bond, U. Deletion of the nuclear exosome component RRP6 leads to continued accumulation of the histone mRNA HTB1 in S-phase of the cell cycle in Saccharomyces cerevisiae. Nucleic Acids Res. 35, 6268–6279 (2007).

97. Jamonnak, N. et al. Yeast Nrd1, Nab3, and Sen1 transcriptome-wide binding maps suggest multiple roles in post-transcriptional RNA processing. RNA 17, 2011–2025 (2011).

98. Grigoriev, I. V. et al. MycoCosm portal: gearing up for 1000 fungal genomes. Nucleic Acids Res. 42, D699–704 (2014).

99. Skrajna, A. et al. U7 snRNP is recruited to histone pre-mRNA in a FLASH-dependent manner by two separate regions of the stem-loop binding protein. RNA 23, 938–951 (2017).

100. Godfrey, A. C. et al. U7 snRNA mutations in Drosophila block histone pre-mRNA processing and disrupt oogenesis. RNA 12, 396–409 (2006).

101. Grigoriev, I. V. et al. Fueling the future with fungal genomics. Mycology 2, 192–209 (2011).

102. Ahrendt, S. R. et al. Leveraging single-cell genomics to expand the fungal tree of life. Nat. Microbiol. 3, 1417–1428 (2018).

