## Supplementary Figure for "Evolution of histone mRNA 3’ end processing machinery in eukaryotes"

#### Supplementary Table S1: Integrated candidates from InterPro, Pfam, CDD and PANTHER

| **Accession** | **Protein** | **Name** |
| --- | --- | --- |
| IPR038294 | SLBP | [SLBP, RNA-binding domain superfamily](https://www.ebi.ac.uk/interpro/entry/InterPro/IPR038294/) |
| IPR026502 | SLBP | [Histone RNA stem-loop-binding protein SLBP1/SLBP2](https://www.ebi.ac.uk/interpro/entry/InterPro/IPR026502/) |
| IPR029344 | SLBP | [Histone RNA hairpin-binding protein, RNA-binding domain](https://www.ebi.ac.uk/interpro/entry/InterPro/IPR029344/) |
| PF15247 | SLBP | [Histone RNA hairpin-binding protein RNA-binding domain](https://www.ebi.ac.uk/interpro/entry/pfam/PF15247/) |
| cd01733 | LSM10 | Like-Sm protein 10 |
| IPR039267 | LSM11 | U7 snRNA-associated Sm-like protein Lsm11 |
| IPR039674 | FLASH | CASP8-associated protein 2 |
| PTHR23253 | eIF4G | EUKARYOTIC TRANSLATION INITIATION FACTOR 4 GAMMA |

#### Supplementary Table S2: NCBI Taxonomy clades with and without SLBP

| **Assignment** | **NCBI Taxonomy Clades** |
| --- | --- |
| 24 clades with SLBP | Bilateria, Cnidaria, Placozoa, Ctenophora, Porifera, Choanoflagellate, Filasterea, Ichthyosporea, Aphelida, Fungi, Amoebozoa, Streptophyta, Prasinodermophyta, Stramenopiles, Alveolata, Rhizaria, Haptista, Metamonada, Discoba, Rhodophyta, Cryptophyceae, Glaucocystophyceae, Eukaryota incertae sedis and Apusozoa |
| 2 clades without SLBP | Provora and Hemimastigophora |

#### Supplementary Table S3: NCBI CDD used to filter out core histones

| **CDD** | **CDD_Shortname** |
| --- | --- |
| CDD:467020 | HFD_H2A |
| CDD:467035 | HFD_H2B |
| CDD:467036 | HFD_H3 |
| CDD:467037 | HFD_H4 |
| CDD:128694 | H4 |
| CDD:128705 | H3 |
| CDD:177669 | H4; Provisional |
| CDD:177733 | H3; Provisional |
| CDD:165721 | H2A; Provisional |
| CDD:177756 | H2A; Provisional |
| CDD:165723 | H2A; Provisional |
| CDD:215080 | H2AX; Provisional |
| CDD:177758 | H2A; Provisional |
| CDD:215081 | H2B; Provisional |
| CDD:165727 | H3; Provisional |
| CDD:215082 | H3; Provisional |
| CDD:165730 | H4; Provisional |
| CDD:185397 | H4; Provisional |
| CDD:185399 | H2A; Provisional |
| CDD:185400 | H3; Provisional |
| CDD:240330 | H2A; Provisional |
| CDD:185642 | H2B; Provisional |
| CDD:355063 | H2B; Provisional |
| CDD:197718 | H2B |
| CDD:197711 | H2A |

#### Supplementary Table S4: NCBI annotations used to exclude core histone candidates

C-terminus of histone H2A

CENP-T histone fold, kinetochore scaffold protein

CENP-T/Histone H4 histone fold domain-containing protein

CENP-T/Histone H4, histone fold

CSE4p Centromere protein that resembles histone H3

Cell cycle-regulated histone H1-binding protein

Centromere kinetochore component CENP-T histone fold

Centromere protein that resembles histone H3

MAG: SIR2 family histone

MAG: histone

MAG: histone 2A

MAG: histone 3

MAG: histone H1

MAG: histone H2A

MAG: histone H2A-1

MAG: histone H2A.1

MAG: histone H2A.V

MAG: histone H2A.Z

MAG: histone H2B

MAG: histone H3

MAG: histone H3-like protein

MAG: histone H3.1

MAG: histone H3.3

MAG: histone H4

MAG: histone MYST family

NASP family histone binding protein

NASP family protein histone binding protein

Sir2 family histone

TPA: histone H2A

TPA: histone H2AZ

TPA: histone H4

TPA_exp: histone H3

Oocyte-specific histone RNA stem-loop-binding protein 2

Zinc finger, H2C2-type, histone UAS binding

Zinc finger, H2C2-type, histone UAS binding protein

cell cycle-regulated histone H1-binding protein

cell cycle-regulated histone h1-binding protein

histone H1-binding

histone H1-binding protein

histone h1-binding

histone h1-binding protein

histone h1-like nucleoprotein HC2 domain-containing protein

methylated histone binding

potential histone binding protein

histone H4-like TAF

histone acetylation-related protein

histone acetyltransferase-like protein esa1

histone binding protein

histone domain-containing protein

histone H3/H4 domain-containing protein

histone H4 replacement-like protein

histone myst family

sir2 family histone

core histone H2A/H2B/H3/H4-domain-containing protein

core histone h2A/H2B/H3/H4 domain-containing protein

Core histone H2A/H2B/H3/H4 family protein

C-terminus of histone H2A

#### Supplementary Table S5: Rfam-curated histone stem-loop sequences not recovered by the custom script

>QBIY01006778.1/511162-511207 QBIY01006778.1 Labeo rohita breed JAYANTI isolate DASCIFA01 scaffold_6824, whole genome shotgun sequence

ccaACAGCUCUUUUAAGAGCCAcccacauuuucaaaaaaauaguuu

......(((((....))))).......................... ( -5.60)

>QBIY01011656.1/1259486-1259531 QBIY01011656.1 Labeo rohita breed JAYANTI isolate DASCIFA01 scaffold_11753, whole genome shotgun sequence

ccaACAGCUCUUUUAAGAGCCAcccacauuuucaaaaaaauaguuu

......(((((....))))).......................... ( -5.60)

>NC_041022.1/23557126-23557171 NC_041022.1 Parambassis ranga chromosome 1, fParRan2.1, whole genome shotgun sequence

CCAAAGGCCCUUUUAAGAGCCACACACUUUCACAUAAAAAGUUUCU

.....(((.((....)).)))....(((((.......))))).... ( -5.00)

>LR778275.1/4248502-4248454 LR778275.1 Coregonus sp. 'balchen' genome assembly, chromosome: 23

CCAAAGGCUCUUUUUUAAAGAGCCACCCACAUUGCAAUAAAAAUAGUAU

.....((((((((...))))))))......................... (-10.70)

>CM031323.1/24030345-24030391 CM031323.1 Anguilla anguilla isolate tag_6206 chromosome 4, whole genome shotgun sequence

CCAAAGGCUCUUUUUAAGAGCcacccacuuuuuaaaaauagcauuau

.....(((((((...)))))))......................... ( -9.80)

>VZRZ01000609.1/10108-10153 VZRZ01000609.1 Climacteris rufus voucher MVMA:354 isolate B10K-DU-029-53 scaffold2977, whole genome shotgun sequence

CCAAAGGCCCUUUUAAGAGCCACCCACAUAAUCUAUAAAGGAGCUU

....(((((((((..(((.............)))..))))).)))) ( -9.00)

>CM016646.2/24690543-24690589 CM016646.2 Camelus dromedarius isolate Drom800 breed African chromosome 20, whole genome shotgun sequence

CAAAAGGCUCUUUUUCAGAGCCACCCAUGUAUCCAAAAAAGGGCUGU

.....((((((.....)))))).(((..............))).... (-11.50)

>QNTS01016779.1/960-915 QNTS01016779.1 Hucho hucho isolate haploid_2015_munich scaffold16780, whole genome shotgun sequence

ccaaaggcuccUUUAAGAGCCACCCACAUAGCAAUAAGAGUGUUCU

.....(((((......)))))...(((............))).... ( -8.40)

>CM014938.1/4693311-4693358 CM014938.1 Perca flavescens isolate YP-PL-M2 chromosome 4, whole genome shotgun sequence

CAAAAGGCUCUUUUUAAGAGCCACCACACAUUCACAAGAGAGCUGUUU

.....(((((((...))))))).....((((((......))).))).. (-10.90)

>NC_052311.1/19205389-19205434 NC_052311.1 Salvelinus namaycush isolate Seneca chromosome 5, SaNama_1.0, whole genome shotgun sequence

ccaaaugcucuuuuaagagccacccACAUUaaaauaagaguauucu

...(((((((((.......................))))))))).. ( -6.70)

>LR584416.1/22890602-22890650 LR584416.1 Salmo trutta genome assembly, chromosome: 3

ccaaaggcucuuuuauuAAGAGCCAcccacauugcaauaagaguauucu

.....(((((((.....)))))))......................... (-10.90)

>NC_027304.1/24725867-24725915 NC_027304.1 Salmo salar isolate Sally breed double haploid chromosome ssa05, ICSASG_v2, whole genome shotgun sequence

CCAAAGGCUCUUUUAUUAAGAGCCACCCACAUUGCAAUAAGAGUAUUCU

.....(((((((.....)))))))......................... (-10.90)

>QNTS01000237.1/17069-17024 QNTS01000237.1 Hucho hucho isolate haploid_2015_munich scaffold237, whole genome shotgun sequence

CCAAAGGCUUUUUUAGGAGCCAUCCACAUUGCAAUAAaaguauucu

.....((((((....))))))......................... ( -6.70)

>LR778283.1/4997760-4997714 LR778283.1 Coregonus sp. 'balchen' genome assembly, chromosome: 31

CCAAAGGCUCUUUCUAAGAGCCACCCACAUUGCAAUAAGAGUAUUUU

.....(((((((...)))))))......................... ( -9.80)

>OV170224.1/18390759-18390713 OV170224.1 Brenthis ino genome assembly, chromosome: 4

CAAACGGCUUUUCUAGAAGCUACUGaacauacuaaaaaauaaauugu

.....((((((....)))))).......................... ( -4.70)

>NW_023010689.1/31941796-31941841 NW_023010689.1 Pantherophis guttatus isolate EG266 unplaced genomic scaffold, UNIGE_PanGut_3.0 Super-Scaffold_11, whole genome shotgun sequence

CCAAAGGCUCUUUUAAAAGCCaccaauaacaaaauaaaaagaauuu

.....((((........))))......................... ( -5.00)

>NC_041231.1/87789159-87789114 NC_041231.1 Physeter catodon isolate SW-GA chromosome 18, ASM283717v5, whole genome shotgun sequence

ACAAAAGCUCUUUUCAGAGCCAucaacauuuucauuuaauaGUUAU

......(((((....))))).......................... ( -5.50)

>SGJD01001421.1/33192-33237 SGJD01001421.1 Balaenoptera physalus isolate FinWhale-01 scaffold1698, whole genome shotgun sequence

ACAAAAGCUCUUUUCAGAGCCaucaacauuuucauuuaauaguuau

......(((((....))))).......................... ( -5.50)

>CACVKT020003233.1/8627-8675 CACVKT020003233.1 Mytilus coruscus strain wild genome assembly, contig: Mco3471, whole genome shotgun sequence

UAAACGGCCCUUUUUUCAGGGCcaccaaauauuucaaaaaagauUCUUU

.....((((((......)))))).......................... (-11.60)

>LR778262.1/33471592-33471547 LR778262.1 Coregonus sp. 'balchen' genome assembly, chromosome: 10

ccaaaggcucuuuuaagUGCCACCCACAUUGCAAUACGAGUAUUCU

.....(((.((....)).)))......................... ( -5.00)

>CACVKT020009619.1/7133-7176 CACVKT020009619.1 Mytilus coruscus strain wild genome assembly, contig: Mco9219, whole genome shotgun sequence

CAAACGGCCCUUUCAGGGCCACCAAUAUUUUCAAAAAGAGUUUA

.....((((((...))))))........................ (-11.00)

>CM007746.2/18725426-18725380 CM007746.2 Oncorhynchus kisutch isolate 150728-3 linkage group LG29, whole genome shotgun sequence

CCAAAGGCUCUUUUUAAGAGCCAUCCACAUUGCAAUAAGAGUAUUCU

.....(((((((...)))))))......................... ( -9.80)

>CM009234.1/16150378-16150332 CM009234.1 Oncorhynchus tshawytscha isolate Chilliwack Hatchery Individual DE9421 chromosome 33, whole genome shotgun sequence

CCAAAGGCUCUUUUUAAGAGCCAUCCACAUUGCAAUAAGAGUAUUCU

.....(((((((...)))))))......................... ( -9.80)

>LR584441.1/68409565-68409519 LR584441.1 Salmo trutta genome assembly, chromosome: 12

ccaaaggcucuuuuuaAGAGCCauccacauugcaauaagaguauucu

.....(((((((...)))))))......................... ( -9.80)

>NC_027310.1/64523124-64523078 NC_027310.1 Salmo salar isolate Sally breed double haploid chromosome ssa11, ICSASG_v2, whole genome shotgun sequence

CCAAAGGCUCUUUUUAAGAGCCAUCCACAUUGCAAUAAGAGUAUUCU

.....(((((((...)))))))......................... ( -9.80)

>QNTS01010442.1/20739-20785 QNTS01010442.1 *Hucho hucho* isolate haploid_2015_munich scaffold10443, whole genome shotgun sequence

ccaaaggcucuuuuuaagagccauccacauugcaauaagaguauucu

.....(((((((...)))))))......................... ( -9.80)

>LR778258.1/9395013-9394968 LR778258.1 Coregonus sp. 'balchen' genome assembly, chromosome: 6

CCAAAAgcucuuuuaagagccacccACAUUGCAAUUAGAGUAUUCU

......(((((....))))).......................... ( -5.60)

>NC_052309.1/27856336-27856290 NC_052309.1 *Salvelinus namaycush* isolate Seneca chromosome 3, SaNama_1.0, whole genome shotgun sequence

ccaaaggcucuuuuuaagagccauccacauuGUAAUAacaguauucu

.....(((((((...)))))))......(((((....)))))..... (-11.30)

>WBNM01011695.1/6896-6941 WBNM01011695.1 Ramphastos sulfuratus voucher USNM:607736 isolate B10K-DU-001-30 scaffold2627, whole genome shotgun sequence

CCAAAGGCCCUUUUAAGAGCCACCCACAUACUCUUUGAAAGAGCUU

....((((.((((((((((...........))).))))))).)))) (-10.00)

>NC_047091.1/86244879-86244834 NC_047091.1 Geotrypetes seraphini chromosome 8, aGeoSer1.1, whole genome shotgun sequence

CCAAAGACUCUUUUAAGAGCCACCCACGCUUUCAAAGAAGAGCUGU

....((.(((((((.(((((.......)))))...))))))))).. (-10.50)

>NW_022990010.1/657552-657597 NW_022990010.1 *Gymnodraco acuticeps* unplaced genomic scaffold, fGymAcu1.1, whole genome shotgun sequence

acaacggcucuuuuaauagccaccuacaguuucaaaaaguugcaau

.....((((........))))......................... ( -4.80)

>NW_022990010.1/735612-735657 NW_022990010.1 Gymnodraco acuticeps unplaced genomic scaffold, fGymAcu1.1, whole genome shotgun sequence

acaacggcucuuuuaauagccaccuacaguuucaaaaaguugcaau

.....((((........))))......................... ( -4.80)

>LR584409.1/1912419-1912464 LR584409.1 Salmo trutta genome assembly, chromosome: 9

ccaaagacucuuuuaagagccauccaCAUuacaauaagaguauucu

......((((((.......................))))))..... ( -3.80)

>CM014940.1/34994995-34995040 CM014940.1 Perca flavescens isolate YP-PL-M2 chromosome 6, whole genome shotgun sequence

ACAAAGGCUCUUUUAAAAGCCACUCUCUUAUUCAUUAAAAAGCUGU

.....((((........))))......................... ( -5.00)

>CACVKT020002343.1/33915-33960 CACVKT020002343.1 Mytilus coruscus strain wild genome assembly, contig: Mco2670, whole genome shotgun sequence

AAAACGGCCCUUUCAGGGCcaccaaauauuucaaaaaagauUCUUU

.....((((((...)))))).......................... (-11.00)

>ABAV01048734.1/586-541 Nematostella vectensis strain CH2 x CH6 NEMVEscaffold_4894_Cont48734, whole genome shotgun sequence.

AAAACGGCUCUUUUAGGAGCUACCACAUGCAAAAAAGUCAAUAGCC

.....((((((....))))))......................... ( -7.00)

#### Supplementary Table S6: Interpro HMMs used to screen and exclude unrelated nucleotidyltransferases

| **Accession** | **Name** |
| --- | --- |
| PF17797 | RL domain |
| PF16367 | RNA recognition motif |
| PF19088 | TUTase nucleotydyltransferase domain |
| PF18528 | RNA editing 3’ terminal uridylyl transferase 2 middle domain |
| PF25420 | ZN292-like C2H2 zinc finger |
| PF12874 | Zinc-finger of C2H2 type |
| PF13912 | C2H2-type zinc finger |
| PF00096 | Zinc finger, C2H2 type |
| SSF54928 | RNA-binding domain, RBD |
| PF00076 | RNA recognition motif |
| PTHR23270 | Programmed cell death protein 11 pre-rRNA processing proteins RRP5 |
| SSF48371 | ARM repeat |
| PF00153 | Mitochondrial carrier protein |
| PF00098 | Zinc knuckle |
| PF09770 | Topoisomerase II-associated protein PAT1 |

#### Supplementary Table S7: Fungal RNA-seq datasets surveyed for histone transcript termini

| **Species** | **SRA** | **Instrument** | **Selection** | **Source** | **Construction protocol** | **SLBP detected** | **SLBP expression (mean depth)** |
| --- | --- | --- | --- | --- | --- | --- | --- |
| **Chytridiomycota** | | | | |  |  |  |
| *Spizellomyces punctatus* DAOM BR117 | SRR343043 | Illumina HiSeq 2000 | cDNA | Transcriptomic | / | Yes | Expressed  (66.5x) |
| *Anaeromyces robustus* | SRR4063399 | Illumina HiSeq 2000 | RT-PCR | Transcriptomic | Regular (RNA) | Yes | Expressed  (15.3x) |
| **Blastocladiomycota** | | | | | |  |  |
| *Allomyces macrogynus* ATCC 38327 | SRR343045 | Illumina HiSeq 2000 | cDNA | Transcriptomic | / | Yes | Expressed  (6.3x) |
| *Catenaria anguillulae* PL171 | SRR424218 | Illumina HiSeq 2000 | cDNA | Transcriptomic | / | Yes | Expressed  (17.2x) |
| **Mucoromycota** | | | | | |  |  |
| Mucor circinelloides | SRR26987983 | Illumina NovaSeq 6000 | cDNA | Transcriptomic | / | No | / |
| *Rhizopus microsporus* | SRR31834734 | Illumina NovaSeq 6000 | cDNA | Transcriptomic | Total RNA was extracted using the RNeasy Plant Mini Kit (Qiagen) and treated with DNase (Sigma, On-Column DNaseI treatment set) NEBNext® Ultra™ Directional RNA Library Prep Kit | No | / |
| **Zoopagomycota** | | | | | |  |  |
| *Entomophthora muscae* | ERR13804685 | DNBSEQ-G400 | Oligo-dT | Transcriptomic | / | No | / |

#### Supplementary Table S8: RNA-seq datasets used to assess expression of histone 3’ end -processing factors and core histone genes in Spizellomyces punctatus DAOM BR117

| **Sample** | **SRA** | **Instrument** | **Selection** | **Source** | **Construction protocol** |
| --- | --- | --- | --- | --- | --- |
| **Study:** Adenine DNA methylation associated with transcriptionally permissive chromatin is widespread across eukaryotes | | | | | |
| *Spun_Multinucleated_rep1* | SRR33446575 | Illumina NovaSeq 6000 | cDNA | Transcriptomic | Fixed cell pellets are treated with 2 ul of 3%SDS, to achieve a final concentration of 0.5% SDS. Cells are thoroughly re-suspended and lysed on ice for 10 min. Lysates are transferred into 0.1 ml Bioruptor Microtubes (Diagenode C30010015, Liège, Belgium) and the chromatin was sheared using an NGS Bioruptor Sonicator (Diagenode) at High Intensity and cycles of 30'' ON/30'' OFF. RNA extracted with NEB Monarch RNA kit ChIP-seq / mRNA-seq |
| *Spun_Multinucleated_rep2* | SRR33446574 | Illumina NovaSeq 6000 | cDNA | Transcriptomic |  |
| *Spun_Zoospores_rep1* | SRR33446573 | Illumina NovaSeq 6000 | cDNA | Transcriptomic |  |
| *Spun_Zoospores_rep2* | SRR33446576 | Illumina NovaSeq 6000 | cDNA | Transcriptomic |  |

#### Supplementary Table S9: Candidate processing factors and histone genes used for expression analysis

| **NCBI Accession** | **Protein** | **NCBI Definition** |
| --- | --- | --- |
| XP_016607336.1 | SLBP | uncharacterized protein SPPG_05550 |
| XP_016605801.1 | ncPAP | uncharacterized protein SPPG_06761 |
| XM_016750633.1 | SL-positive, PAS-negative H2A | Spizellomyces punctatus DAOM BR117 histone H2A (SPPG_02344), mRNA |
| XP_016611334.1 | H2A | histone H2A |
| XP_016611335.1 | H2B | histone H2B |
| XP_016607979.1 | H2A | Histone H2A |
| XP_016605157.1 | H2A | histone H2A |
| XP_016604090.1 | H3 | histone H3 |
| XP_016604091.1 | H4 | histone H4 |
| XP_016608606.1 | H4 | histone H4 |

#### Supplementary Table S10: Median histone stem-loop distances and mean conservation scores across major clades

| **Clade** | **H2A** | **H2B** | **H3** | **H4** | **Mean conservation** |
| --- | --- | --- | --- | --- | --- |
| *Porifera* | 30.0 | 38.0 | 30.0 | 33.0 | 12.6 |
| *Ctenophora* | 20.0 |  | 33.0 | 50.0 | 12 |
| *Cnidaria* | 23.0 | 24.0 | 23.0 | 32.0 | 13.1 |
| *Placozoa* | 48.0 | 58.0 | 51.0 | 66.0 | 12.9 |
| *Bilateria* | 32.0 | 28.0 | 29.0 | 31.0 | 13.9 |
| *Ichthyosporea* | 78.0 | 61.0 | 84.0 | 64.0 | 9.3 |
| *Discoba* | 56.0 | 99.0 | 89.0 | 208.0 | 5.7 |
| *Metamonada* | 54.0 | 44.0 | 65.0 | 80.0 | 5.7 |
| *Amoebozoa* | 114.0 | 106.0 | 62.0 | 96.0 | 7.9 |
| *Rhizaria* | 23.0 | 25.0 |  |  | 9.0 |
| *Stramenopiles* | 105.0 | 72.0 | 66.0 | 114.0 | 7.9 |
| *Alveolata* | 199.0 | 114.0 | 159.0 | 132.0 | 7.9 |
| *Prasinodermophyta* |  |  |  |  | 0.0 |
| *Chlorophyta* | 59.0 | 54.0 | 57.0 | 61.0 | 8.4 |
| *Klebsormidiophyceae* | 21.0 | 39.0 |  | 70.0 | 6.3 |
| *Bryophyta* | 37.0 | 34.0 | 91.0 | 40.0 | 8.0 |
| *Acrogymnospermae* | 43.0 | 37.0 | 37.0 | 37.0 | 7.2 |
| *eudicotyledons* | 117.0 | 141.0 | 126.0 | 110.0 | 5.0 |
| *Liliopsida* | 114.0 | 106.0 | 135.0 | 48.0 | 4.2 |
| *Microsporidia* | 18.0 | 256.0 | 188.0 | 110.0 | 9.0 |
| *Cryptomycota* | 26.0 | 26.0 | 45.0 | 27.0 | 10.1 |
| *Chytridiomycota* | 81.0 | 114.0 | 84.0 | 93.0 | 7.6 |
| *Blastocladiomycota* | 90.0 | 110.0 | 129.0 | 102.0 | 11.9 |
| *Zoopagomycota* | 182.0 | 48.0 | 184.0 | 253.0 | 5.0 |
| *Mucoromycota* | 142.0 | 109.0 | 182.0 | 133.0 | 5.3 |
| *Ascomycota* | 141.0 | 133.0 | 194.0 | 177.0 | 5.1 |
| *Basidiomycota* | 178.0 | 163.0 | 168.0 | 171.0 | 5.0 |

#### Supplementary Figure S1: Phylogenetic tree of the SLBP RNA-binding domain

Maximum-likelihood phylogenetic tree of 5,926 SLBP RBD sequences across eukaryotic clades. Strip colors indicate major taxonomic groups according to the NCBI taxonomy database.

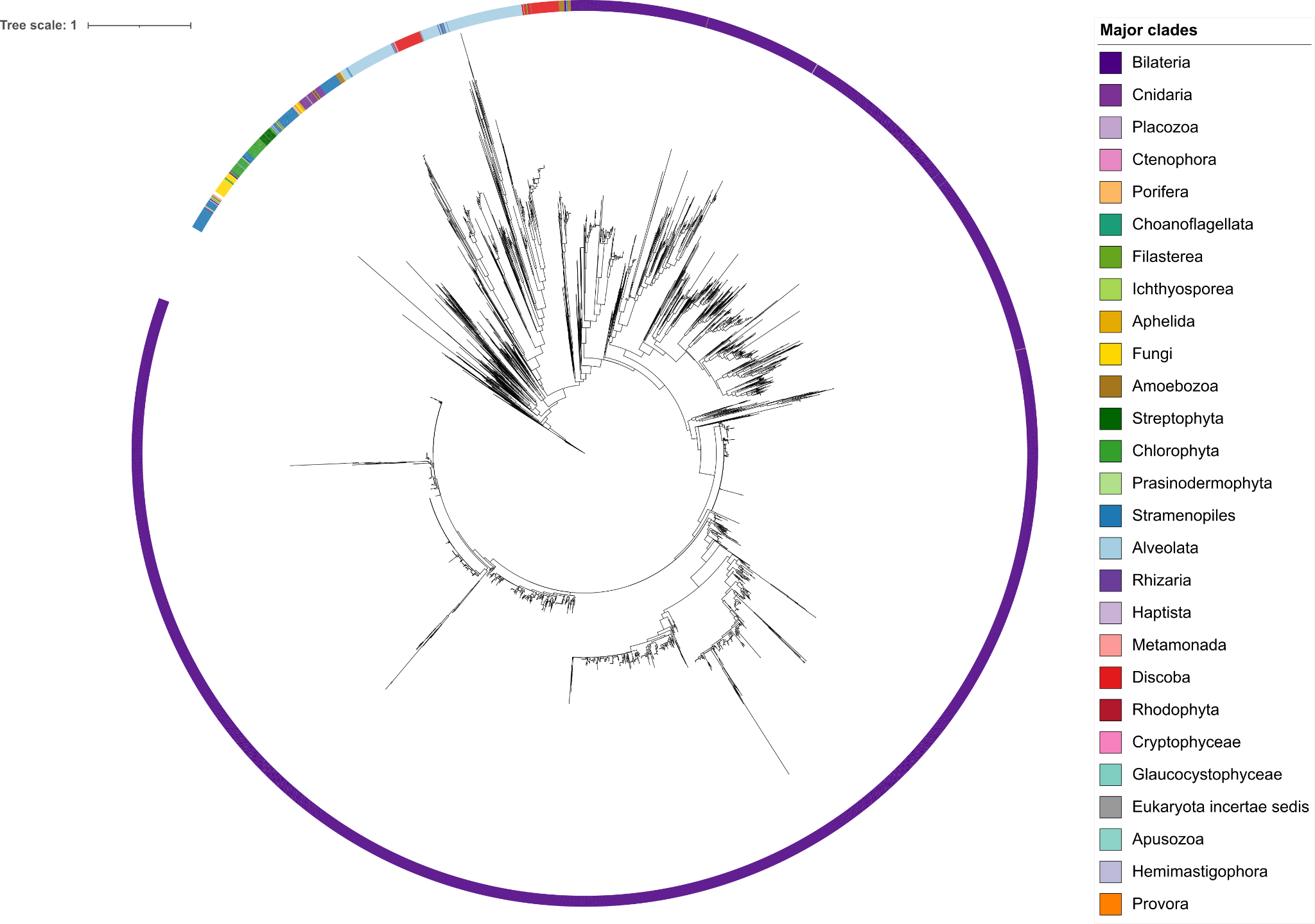

#### Supplementary Figure S2: Public sources of SLBP ortholog annotation across eukaryotes

Phylogenetic distribution of SLBP across all eukaryotic clades, inferred from BLASTP, InterPro, OrthoDB, and TBLASTN searches. Taxa were assigned using NCBI TaxIDs. Coloured circles denote the sources of SLBP in the target clades.

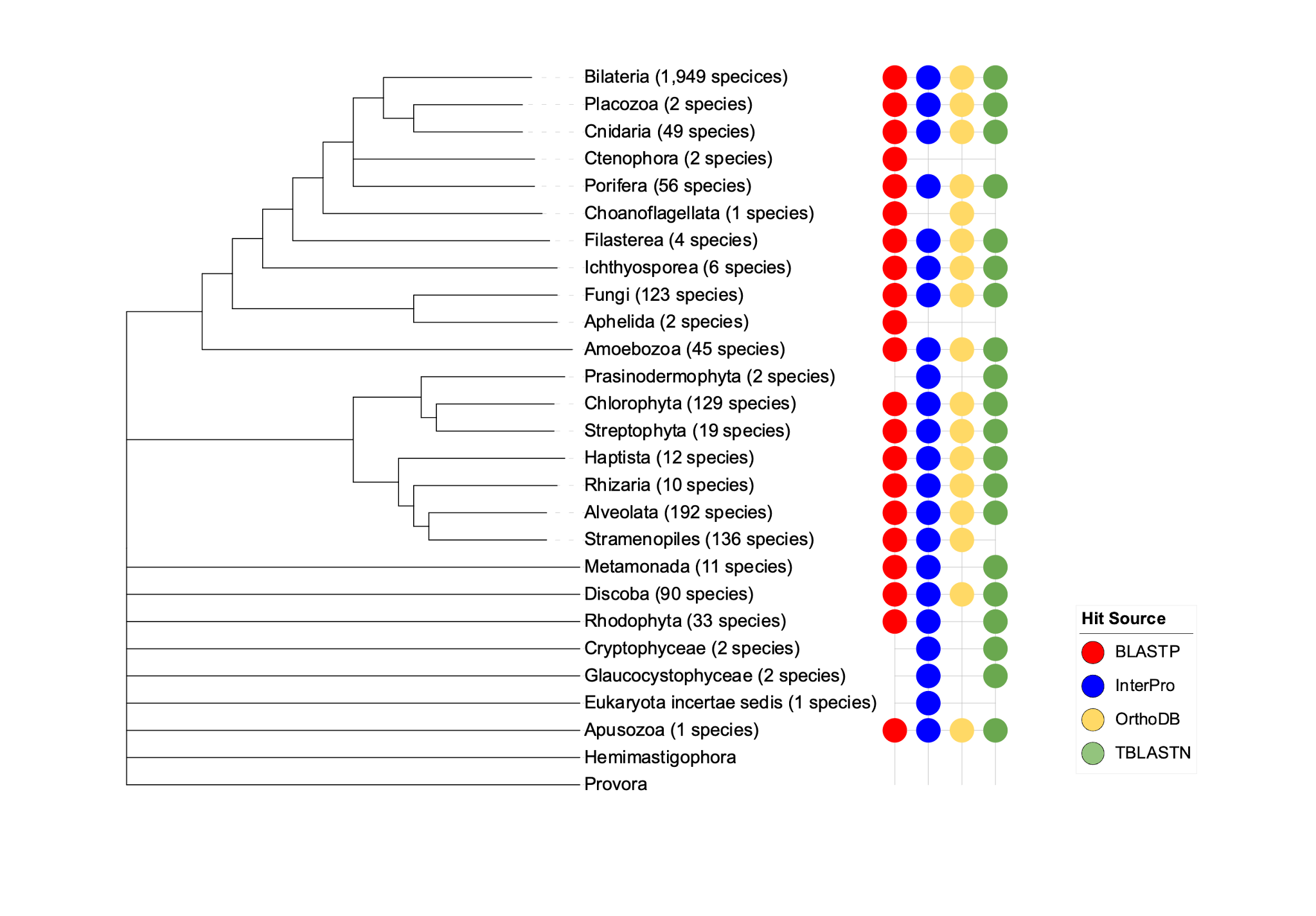

#### Supplementary Figure S3: Sequence conservation of the SLBP SLIP1-interacting motif

Sequence logo of the SLIP1-interacting motif in SLBP, generated using MEME from aligned SLBP protein sequences. The x-axis indicates amino acid position within the motif, and the y-axis shows information content (bits). Letter height is proportional to residue conservation at each position.

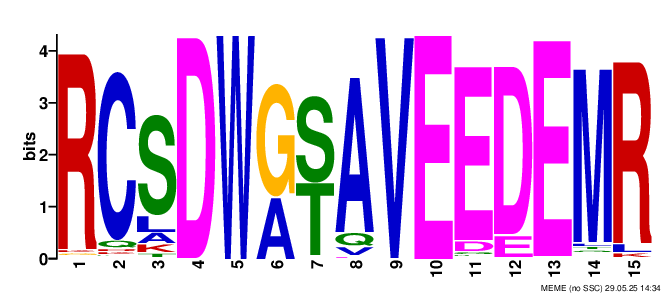

#### Supplementary Figure S4: Phylogeny of SLIP1-like MIF4G-domain proteins

ML tree of short MIF4G-domain proteins used to validate SLIP1 homologs. Sequences (< 300 aa) were filtered to exclude long eIF4G/CTIF-like proteins, retained based on stronger support for PTHR23254 than PTHR23253. Proteins were aligned with MAFFT, the phylogeny was constructed with FastTree annotated using iTOL.

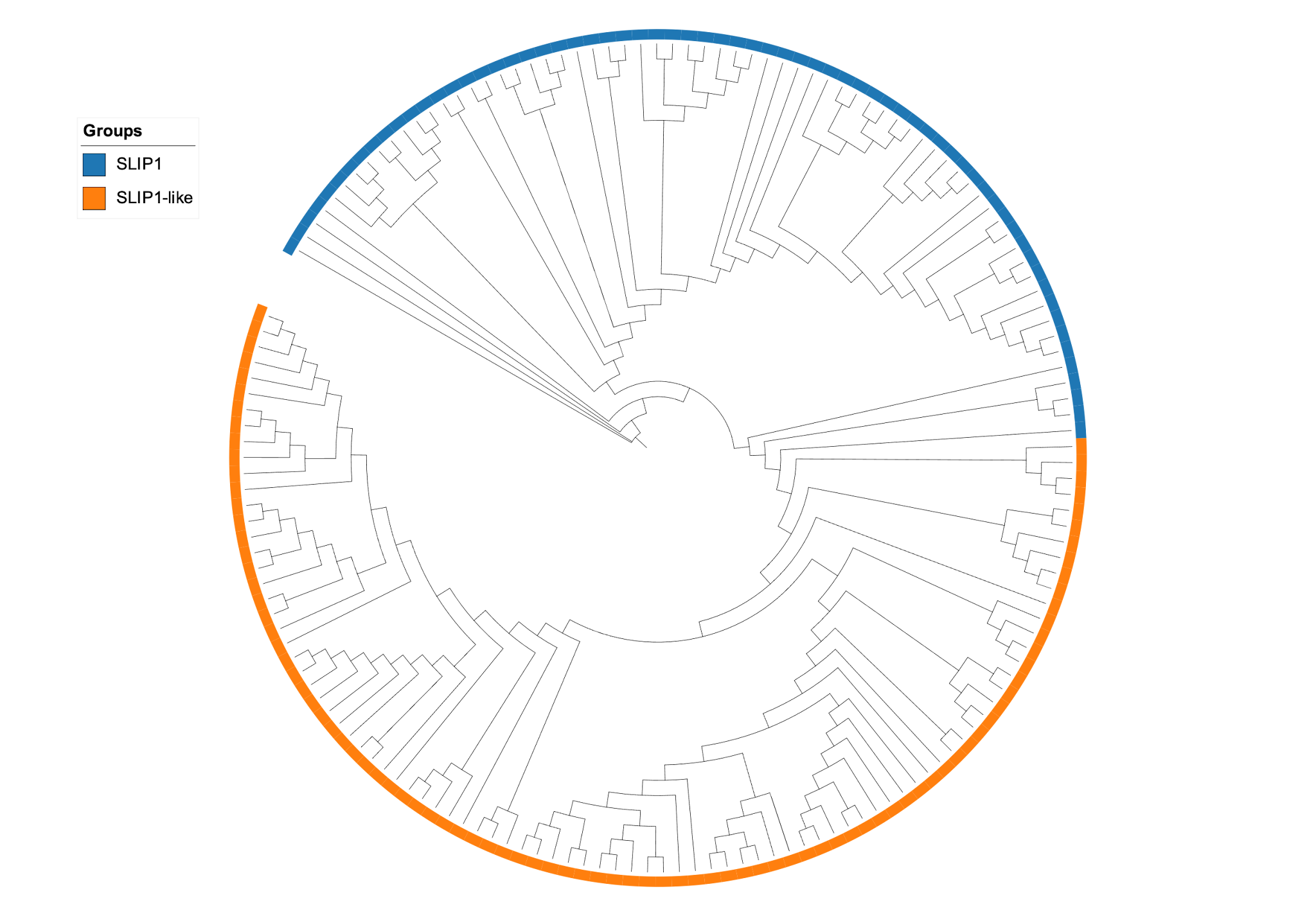

#### Supplementary Figure S5: Sequence similarity between *Takifugu rubripes* FLASH-associated domain query and translated genomic region in *Rozella allomycis* CSF55

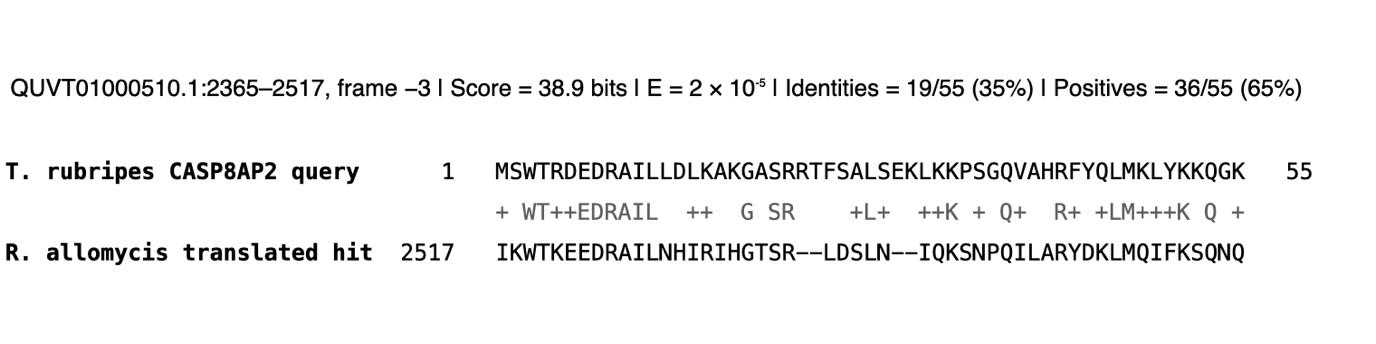

**Supplementary Figure S6: Identification and secondary-structure validation of a U7-like snRNA in Rozella allomycis.**

A U7-like snRNA was detected in the *R. allomycis* SCF55 genome (Accession: ML007894.a) by covariance model search using the Rfam model (RF00066), with a significant hit (e-value = 8.6e-5). The predicted candidate is divergent from canonical U7 snRNAs, but retains key residues within the Sm-binding sites as described in Kolev & Steitz 2005. The secondary structure was predicted and visualized using RNAfold.

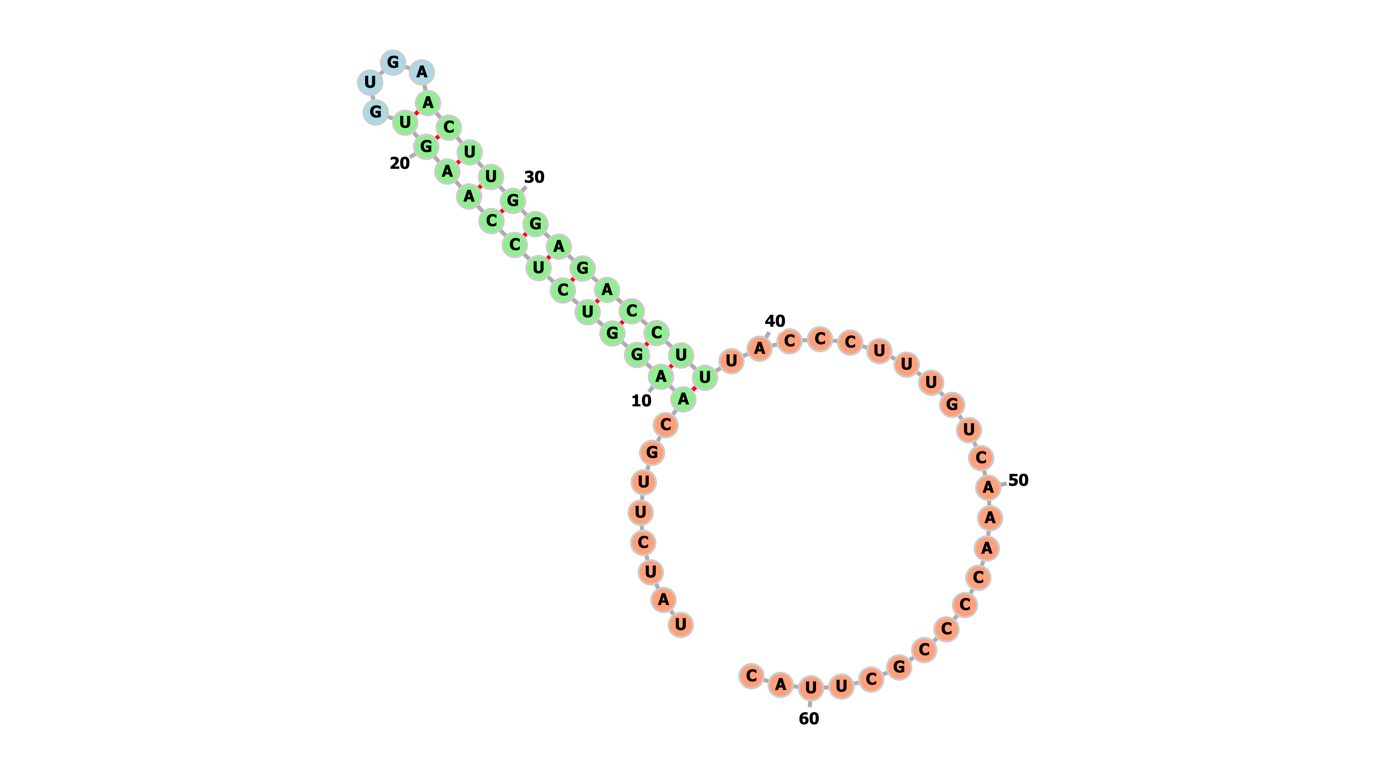

**Supplementary Figure S7A: Similarity network of the 50 most frequent metazoan histone stem loops.** Each node represents one of the top 50 SL sequences, labeled by its index and predicted hairpin fold free energy (ΔG; kcal/mol). The network layout (Kamada-Kawai) places more similar SLs closer together. Node size is normalized by frequency.

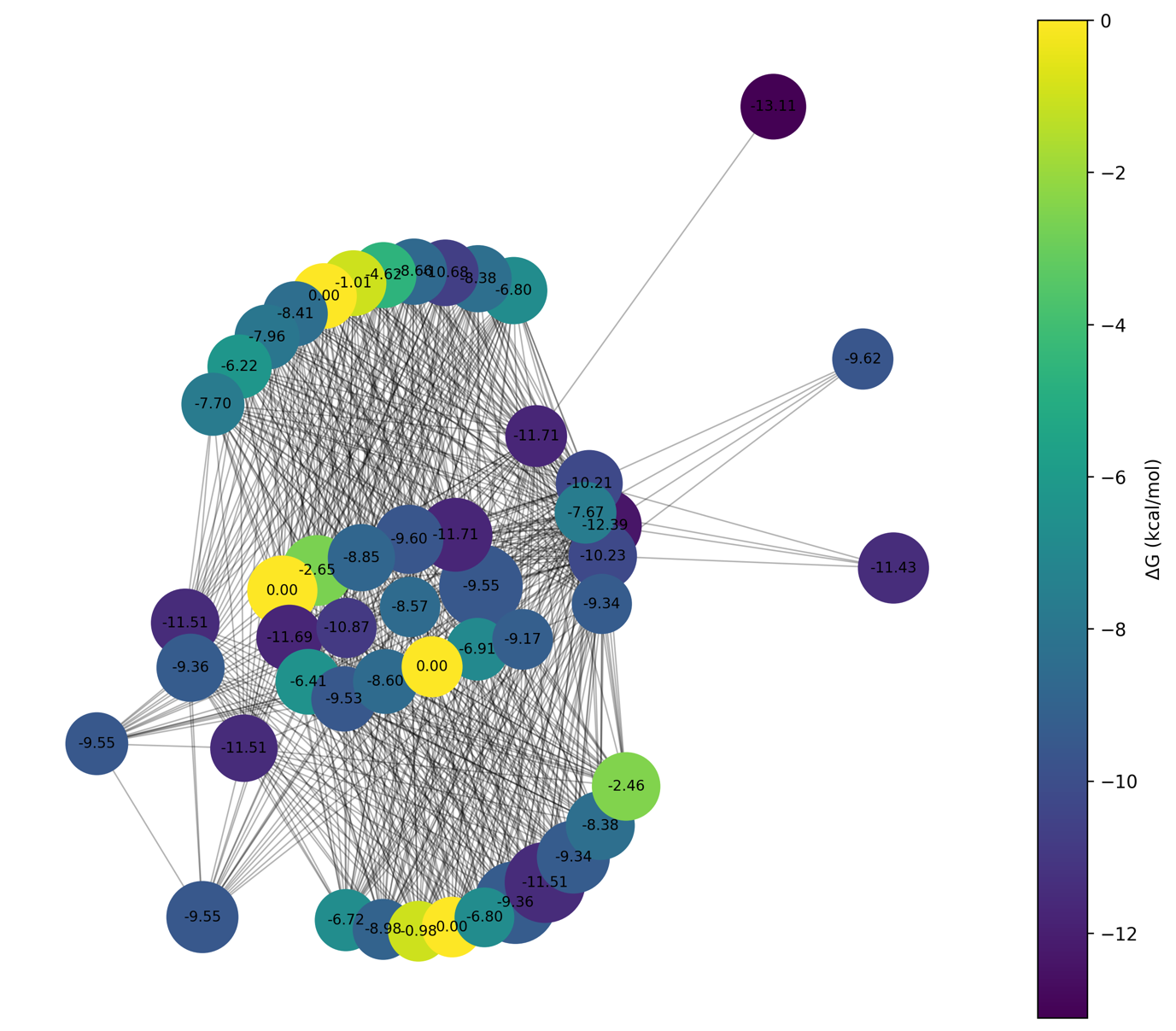

#### Supplementary Figure S7B: Similarity network of the 50 most frequent fungal histone stem loops

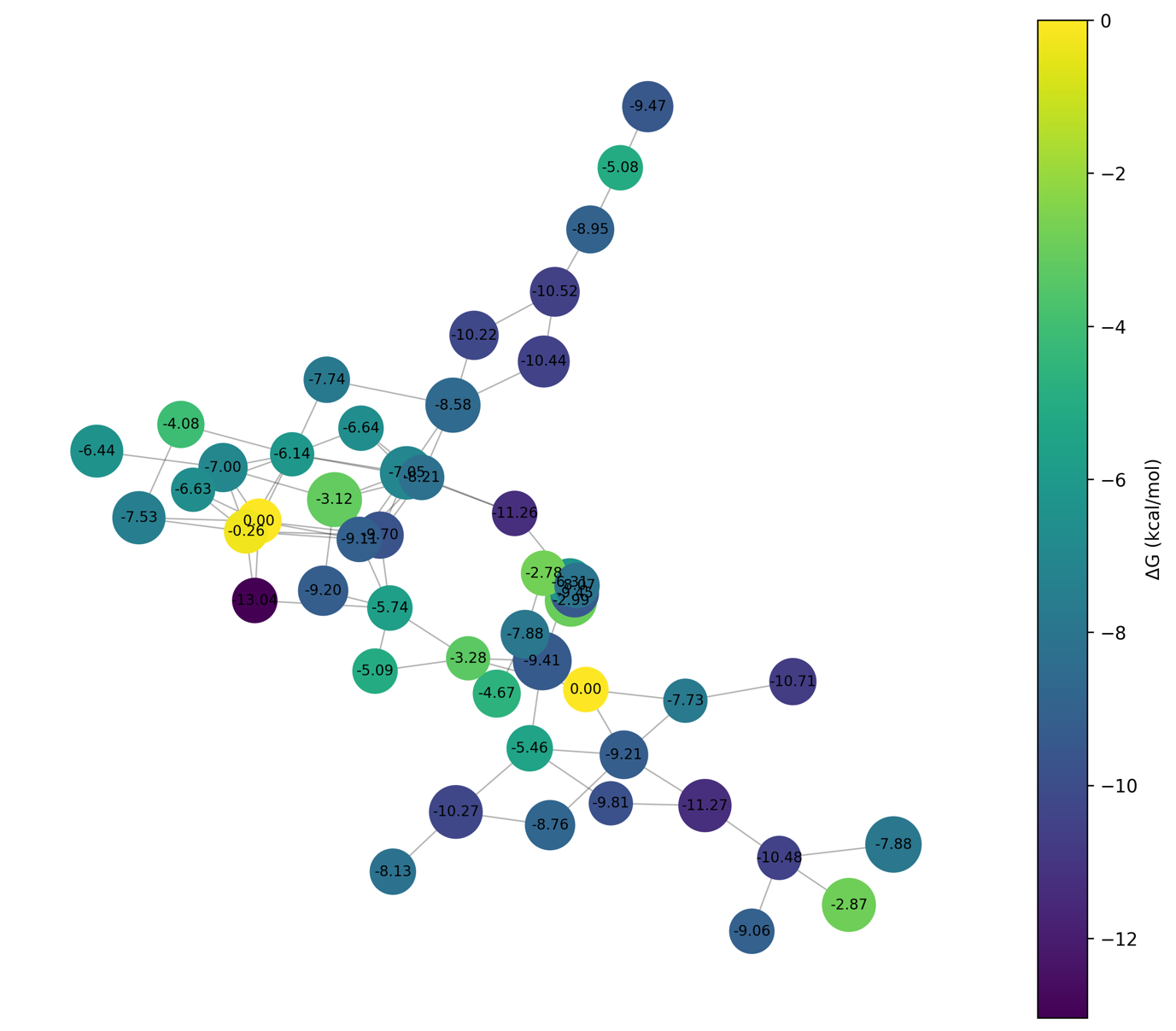

#### Supplementary Figure S7C: Similarity network of the 50 most frequent protozoan histone stem loops

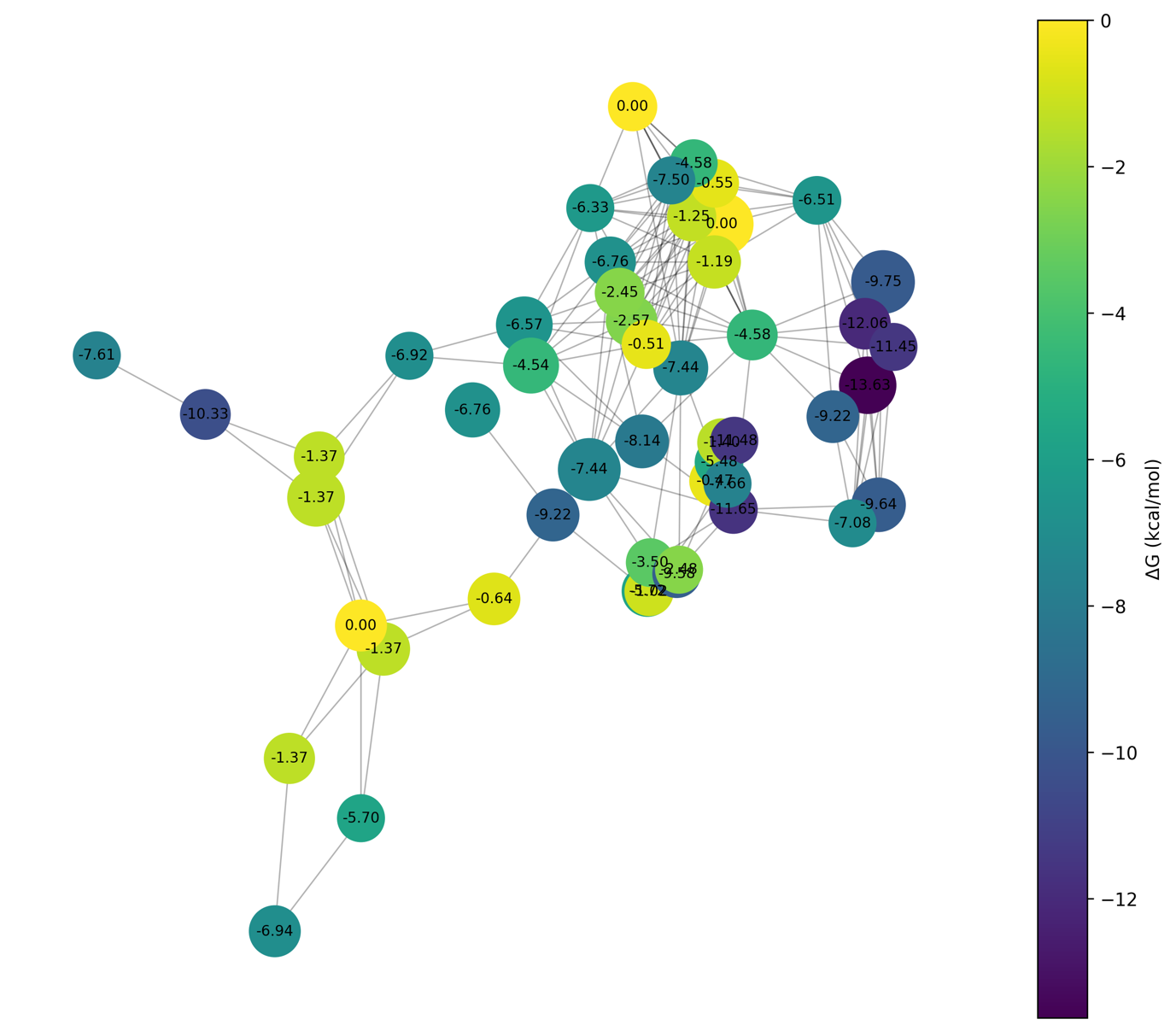

#### Supplementary Figure S7D: Similarity network of the 50 most frequent plant histone stem loops

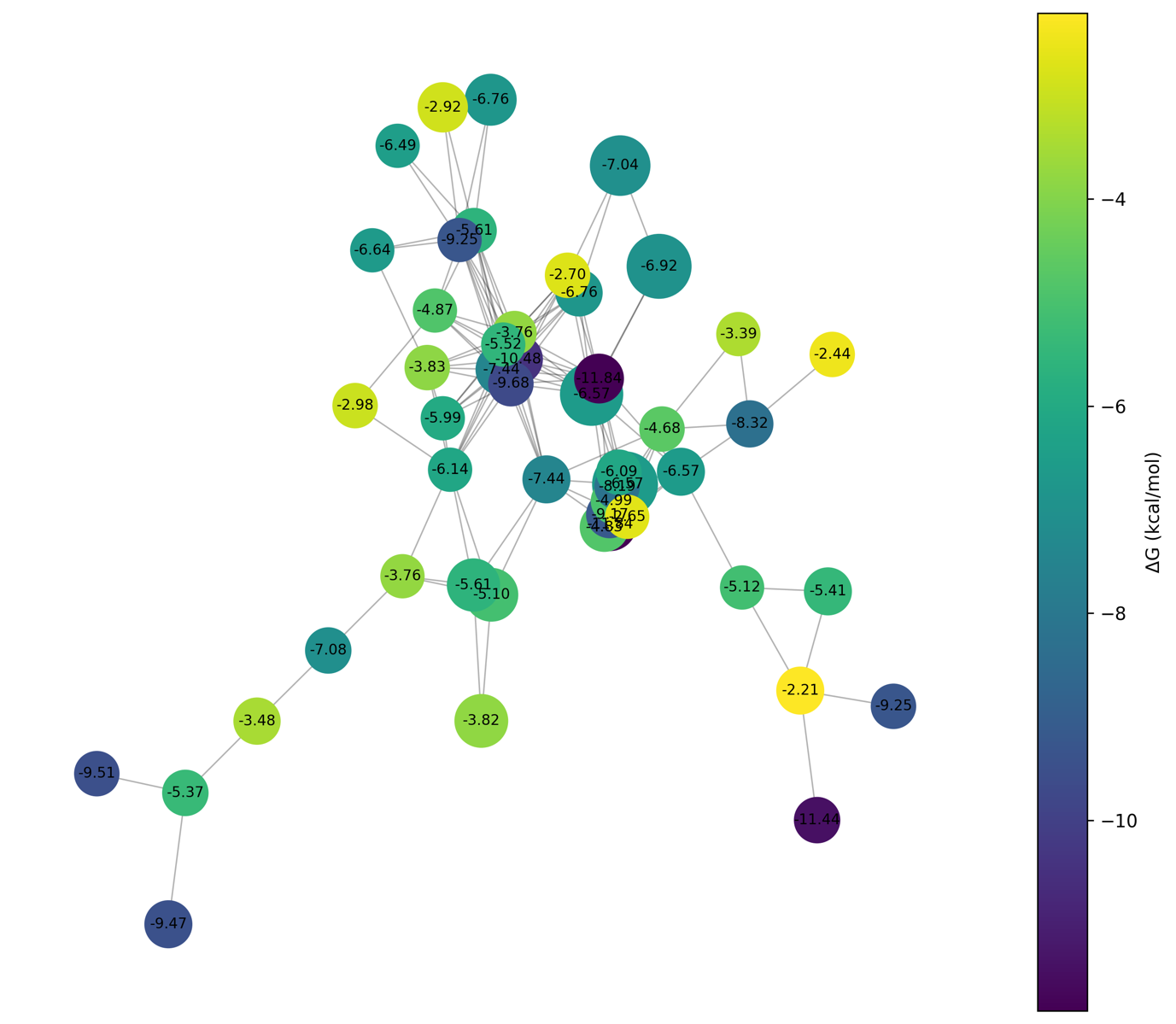

**Supplementary Figure S8: Public mRNA-seq datasets support predicted locus-level 3′-end processing patterns.** Species are grouped by higher-level fungal clade. For each histone locus, the schematic shows the predicted 3′-end features, including the stem-loop (SL), polyadenylation signal (PAS; including weak PAS where indicated), and the poly(A) site inferred from mapped mRNA-seq reads with poly(A) tails. Distances between features are indicated in nucleotides (nt). Histone type and corresponding protein accession are shown at right.

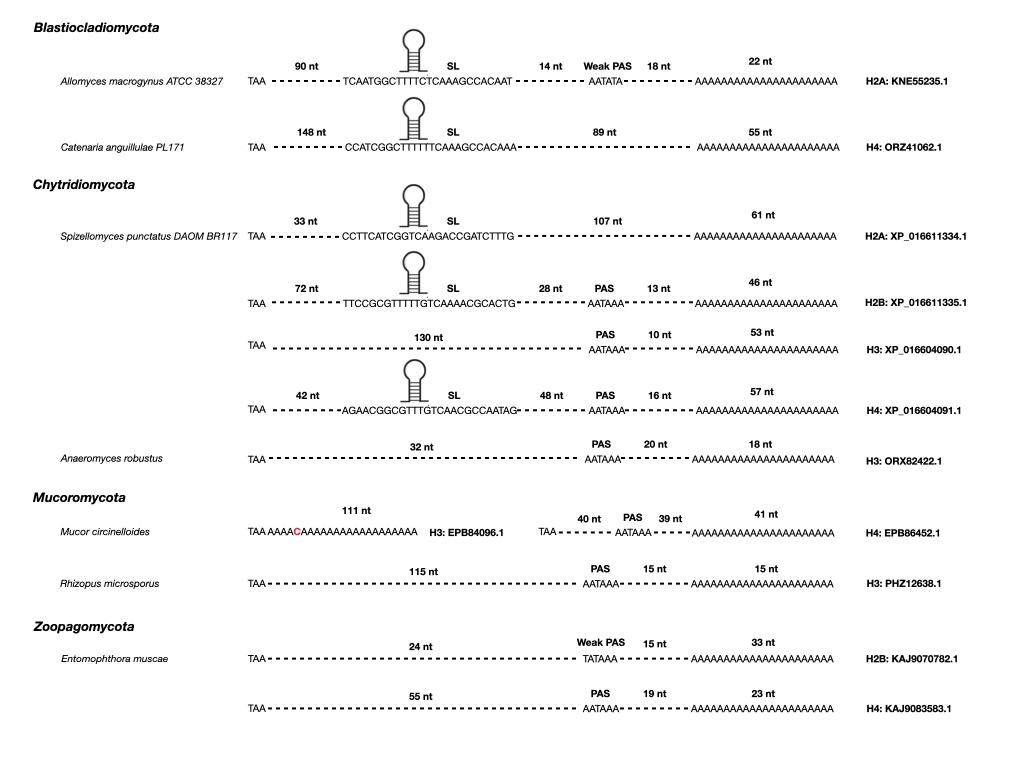

**Supplementary Figure S9:** **Expression of candidate SLBP-associated processing factors and core histones in *Spizellomyces punctatus.*** Public strain-specific mRNA-seq data (accession: PRJNA983123) from *Spizellomyces punctatus* zoospores (Zoo) and multinucleated (Mult) show high expression of candidate ncPAP and SLBP together with core histones in the multinucleated stage. Counts per million (CPM) are calculated as target primary mapped reads divided by total primary mapped reads. Multiplied by 1 million after excluding unmapped, secondary, and supplementary alignments using samtools. The PAS-lacking H2A was kept separate from other H2A.

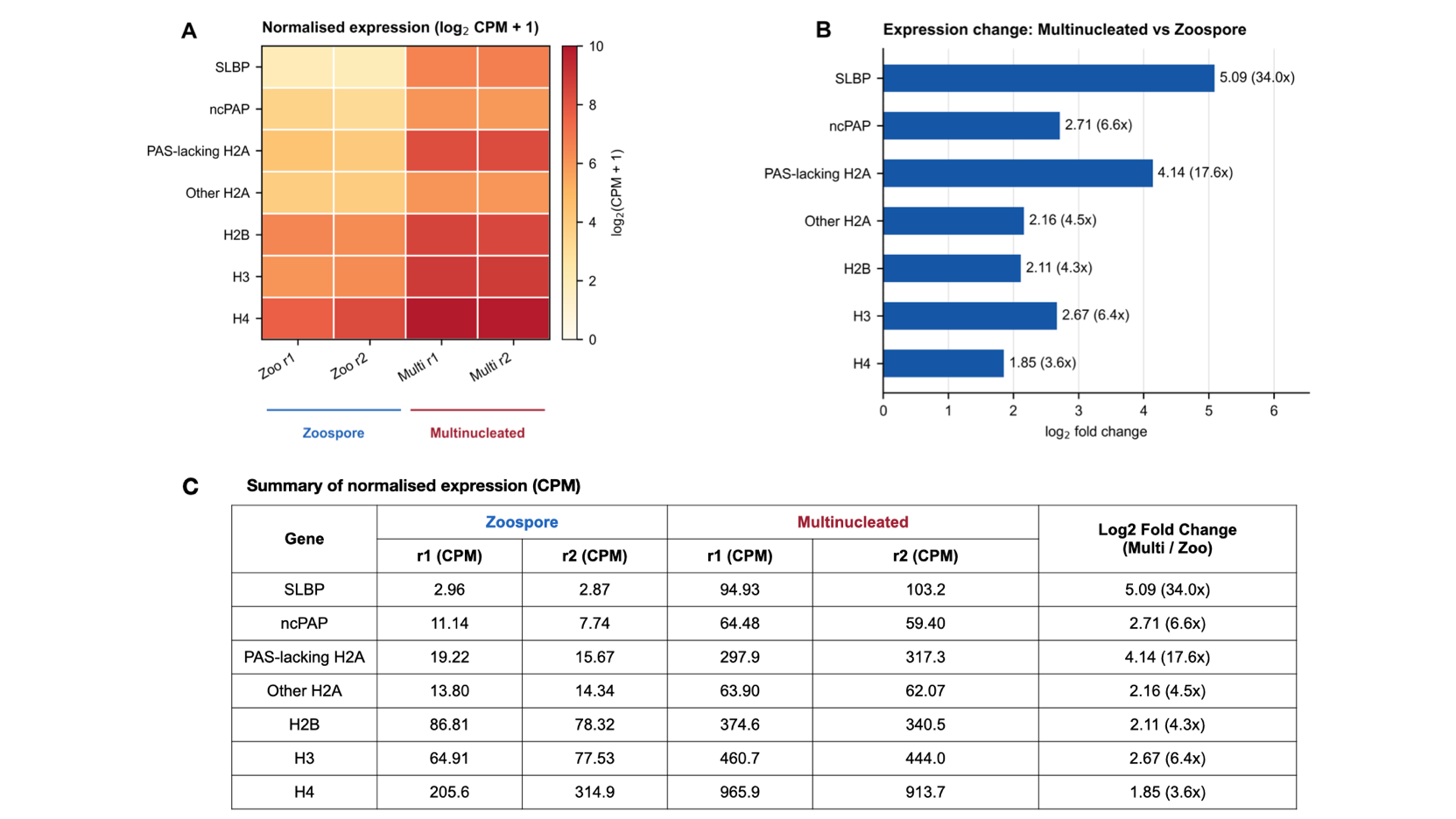
